# Structural Insights into Bromodomain-Containing Complexes from *Trypanosoma cruzi* Revealed by Proximity Labeling and Stoichiometric Space Exploration

**DOI:** 10.64898/2026.03.22.713544

**Authors:** Elvio Rodríguez Araya, Gonzalo Martínez Peralta, Victoria Alonso, Esteban Serra

## Abstract

*Trypanosoma cruzi* is the causative agent of Chagas disease, a neglected illness with outdated treatments. Bromodomain factors (BDFs) are essential proteins that recognize acetylated lysines and have strong therapeutic potential. They form part of epigenetic complexes that regulate chromatin accessibility and, therefore, gene expression. However, little is known about their structure in trypanosomatids. Here, we used a combination of experimental and bioinformatic approaches to infer the stoichiometry and structure of *T. cruzi* bromodomain-containing complexes. By reconstructing the proximity networks of five BDFs using TurboID-directed proximity labeling, we identified highly interconnected components that assemble into the CRKT and NuA4 complexes. Using novel structure prediction strategies that systematically explore the stoichiometric space, we inferred that CRKT assembles into three distinct modules and NuA4 in two, with different degrees of interaction dynamics. The core module of CRKT contains two copies of each component, including BDF3, BDF5, and BDF8, arranged in a subcomplex with central symmetry. The catalytic module of CRKT has three subunits, including the <u>h</u>istone <u>a</u>cetyltransferase 2 (HAT2), while the BET (<u>b</u>romodomain and <u>e</u>xtra-terminal) module has one unit of both BDF4 and BDF1. The catalytic module of NuA4 closely resembles the yeast piccolo-NuA4 module and contains HAT1, while the TINTIN module associates with the catalytic module via the C-terminal domain of BDF6. These insights shed light on the structure and composition of epigenetic complexes in trypanosomatids, opening new avenues for rational drug design aimed at disrupting their function.

## Introduction

*Trypanosoma cruzi* is the protozoan that causes Chagas disease, which affects more than 7 million people worldwide [1]. Developed more than 60 years ago, only two drugs are currently available for its treatment. Both have limited effectiveness during the chronic stage of the disease and usually cause adverse effects that lead to discontinuation of therapy [2]. In addition to the existence of resistant strains, the efficacy of these drugs varies between strains and hosts, even in the acute stage of the disease. Taken together, these disadvantages point the need to develop alternative therapies for this neglected disease [3].

Within this context, trypanosomatid bromodomains (BDs) emerge as potential therapeutic targets [4–6]. With approximately 100 amino acids in length, BDs have four α-helices arranged to form a recognition pocket for acetylated lysines [7]. The specific function of BD containing proteins, known as bromodomain factors (BDFs), is dictated by the BD’s specificity for acetylated lysines, its subcellular localization, the presence of other domains in its architecture, and the proteins with which they interact [8]. Typically, BDFs are part of multiprotein complexes, where the recognition of acetylation signals by the BDs contributes to the complex’s function. In the nucleus, BD containing complexes often contain <u>h</u>istone <u>a</u>cetyltransferase (HAT) enzymes, which acetylate lysine residues in histones; and <u>h</u>istone <u>d</u>e<u>ac</u>etylases (HDACs), which remove these post-translational modifications (PTMs), along with other components related to chromatin remodeling [9]. Besides interacting physically through protein-protein interactions (PPIs), BDFs, HATs, and HDACs interact functionally by recognizing, adding, or removing acetylations on nucleosome lysines, thereby coordinating epigenetic regulatory mechanisms essential for eukaryotic biology [9].

*T. cruzi* has eight BDFs with diverse domain architectures [10]. Some have already been validated as therapeutic targets and the effects of gene deletion/overexpression on the parasite’s life cycle have been studied. BDF2 has been identified as an essential protein, was found in the nucleus at all life stages, it recognizes acetylated lysines of histone H4, and has been detected as an interactor of the variant histone H2B.V [10–12], in consistency with the typical epigenetic role of these proteins. However, BDF1 and BDF3, two members of the BET (Bromo and Extra-Terminal) family that have also been identified as essential components, exhibit extranuclear localization [13]. BDF1 colocalizes with the glycosomal hexokinase and possesses a PTS-2 type peroxisomal localization signal at its N-terminus, suggesting a specific role in glycosomes [14]. In immunoprecipitation experiments, BDF3 is recovered alongside α-tubulin, and their interaction has been demonstrated *in vitro* using acetylated peptides of α-tubulin [15]. Furthermore, BDF3 is consistently detected in the cytoplasm by immunofluorescence [16]. This differential distribution suggests specialized functions outside the nucleus, possibly in the recognition of cytoplasmic acetylated proteins or regulating processes unrelated to transcription. Finally, BDF6 has been identified as an essential component for amastigote growth [17].

On the other hand, interactomic assays of their orthologs have been carried out in related organisms. In *Trypanosoma brucei*, interaction networks constructed from co-immunoprecipitation (co-IP) data of BDFs and other related proteins (such as HATs and HDACs) indicate they form part of multiple complexes with varying degrees of interconnectivity among their components [18, 19]. In *Leishmania mexicana*, a complex essential for transcription, named <u>C</u>onserved <u>R</u>egulators of <u>K</u>inetoplastid <u>T</u>ranscription (CRKT), has been identified using proximity labeling with BDF5 as bait [20]. Identified components include BDF3, BDF5, BDF8, and HAT2, consistent with what has been observed in co-IP interaction networks of *T. brucei* [19]. Furthermore, proximal to BDF5 were detected the factors BDF4 and BDF6 and the histones H2A.Z, H2B.V, H3 and H3.V. However, very little is known about their three-dimensional structural organization, a fundamental requirement for developing targeted therapies that seek to disrupt the normal functioning of the overlaying complexes.

Thanks to advances in protein structure prediction, we recently demonstrated that it is possible to assign a precise structural dimension to these interactomic datasets and experimentally validated the interactions predicted with deep learning methods [21]. In the process, we identified MRG domains as essential elements for the assembly of BDF containing complexes and confirmed the presence of TcTINTIN (trimer independent of NuA4 for transcription interaction), an ortholog of the yeast TINTIN subcomplex, which is part of the complex known as NuA4 [22]. TcTINTIN is composed of the proteins BDF6, MRGx, and MRGBP (MRG Binding Protein), where the presence of the MRG domain of MRGx is essential to stabilize MRGBP and allow its interaction with BDF6. We also identified a ternary complex formed by BDF5, BDF8, and BDF5BP (BDF5 Binding Protein), where the MRG domain of BDF5 interacts simultaneously with BDF8 and BDF5BP, acting as a bridge that connects them.

Beyond these advances, interactomic data for *T. cruzi* are still lacking to confirm the observations made in *T. brucei* and *Leishmania*. Moreover, considering the differences in subcellular localization of BDF1 and BDF3 between *T. cruzi* (extranuclear) [14, 15] and other trypanosomatids (nuclear) [19], fundamental differences between these complexes may exist. In this work, we performed such interactomic experiments using proximity-labeling in *T. cruzi* epimastigotes and BDFs as baits. We reconstructed the proximity networks from the data obtained and identified highly interconnected proteins that assemble the underlying complexes. We verified the subcellular localization of BDFs using indirect immunofluorescence and generated a high-quality nuclear proximity proteome that allowed us to orthogonally validate the observed localizations. Applying the latest methodological advances in protein structural bioinformatics, we systematized structure prediction, analysis, and PPI prediction among the highly interconnected components of the proximity networks, obtaining the three-dimensional structures and the most probable stoichiometries of *T. cruzi* CRKT and NuA4 complexes. To do this, we used MultimerMapper, developed in our laboratory to systematically explore the stoichiometric space using AlphaFold3 predictions. It identifies stable complexes, automatically predicts PPIs, captures conformational changes, and finds the most probable stoichiometries [23–26].

## Results

### Setup of TurboID-based proximity labelling of BDFs in T. cruzi epimastigotes

To resolve the interactomes of *T. cruzi* BDFs, we implemented the proximity labeling method using the highly active biotin ligase TurboID. This technique allows the capture of both stable and transient interactions in live cells, without depending on the physical stability of the complexes during the purification process [27]. Using the pTcTurboID vector, we generated stable *T. cruzi* cell lines expressing fusions to three N-terminal HA tags and TurboID under the control of a tetracycline-regulated promoter [28]. To distinguish specific interactions from nonspecific background noise, two spatial controls were prepared. As a cytoplasmic spatial control, TurboID was fused to GFP. As a nuclear spatial control, TurboID was fused to GFP and three nuclear localization signals (NLS) derived from the simian virus SV40 [29]. By fusing TurboID to an inert protein such as GFP, these controls compensate nonspecific biotinylation resulting from the diffusive process that baits undergo while they are not in their specific locations [30]. As baits, 3xHA-TurboID fusions of BDF2, BDF3, BDF4, BDF5, BDF6, and BDF8 were generated. Epimastigotes of *T. cruzi* Dm28c strain expressing the pLew13 plasmid [31] were transfected with the resulting vectors, and stable clonal populations were selected. For each transfection, the clone with the best regulation profile was selected; that is, those with the lowest possible basal expression and good expression upon induction with tetracycline concentrations that maximize expression (0.5 μg/mL). For all linages, at least one clone with tetracycline-regulated expression was obtained (**Supplementary Fig. 1**). However, after two successful rounds of transfection and cloning, all the BDF2 clones obtained showed some expression without tetracycline. So, we decided to continue using the clone with the best tetracycline response profile.

A critical aspect to optimize was achieving expression levels low enough to preserve the physiological function of the BDFs while still producing sufficient biotinylation to detect proximal proteins [32]. Therefore, we determined the minimum tetracycline concentration at which the system begins to respond and selected the lowest concentration that allows for the detection of biotinylated proteins for each fusion. Based on observations made by Taylor et al. [31], we tested 0, 1, and 5 ng/mL of tetracycline. Since the fetal bovine serum in the LIT medium contains biotin, substrate of TurboID, we also evaluated the need to supplement the medium with it. The results revealed that the serum provides sufficient biotin to activate TurboID-mediated biotinylation, making supplementation unnecessary (**Supplementary Fig. 2**). Furthermore, a gradient in expression levels was observed with increasing tetracycline concentrations, with a proportional increase in detected biotinylated proteins. For each linage, we selected the final tetracycline concentration by approximating the intensity of biotinylated protein bands to that of the 75.2 kDa naturally biotinylated protein 3-methylcrotonyl-CoA carboxylase as reference, identified as the most enriched protein in an enrichment test using the parental strain. For linages expressing the BDF5, BDF6, and BDF8 fusions, expression was observed starting at 1 ng/mL, but we decided to use 2 ng/mL instead of 1 ng/mL because the biotinylation levels were slightly lower than the others. For BDF3 and BDF4, we used 1 ng/mL. In the case of GFP-NLS and BDF2, the small leaky expression was sufficient to reach the desired levels, the latter showing the highest escape when no tetracycline was added.

### TurboID fusions localization by indirect immunofluorescence

Before performing purification tests, we determined the subcellular localization of the fusions by indirect immunofluorescence (IFI) using these tetracycline concentrations. The results of the IFI assays using anti-HA and DAPI are shown in **Fig. 1**. As expected, the GFP fusion localized in the cytoplasm, while the GFP-NLS fusion showed a homogeneously distributed nuclear signal. BDF2 and BDF3 fusions were homogeneously distributed in the nucleus and cytoplasm, respectively, consistent with previous studies. BDF5 and BDF6 fusions showed nuclear signal in all observed cells, mainly in the periphery of the nucleus. On the other hand, BDF4 and BDF8 fusions also showed nuclear signal, with some cells exhibiting some cytoplasmic signal. The nuclear signal of BDF4 was more homogeneous, while BDF8 was observed mainly in the nuclear periphery, similar to BDF5 and BDF6. These observations match the subcellular localizations observed in the TrypTag database for the *T. brucei* orthologs of BDF2, BDF4, BDF5, BDF6, and BDF8 [33], but differ for BDF3, which is detected in the nucleus.

**Fig. 1:**
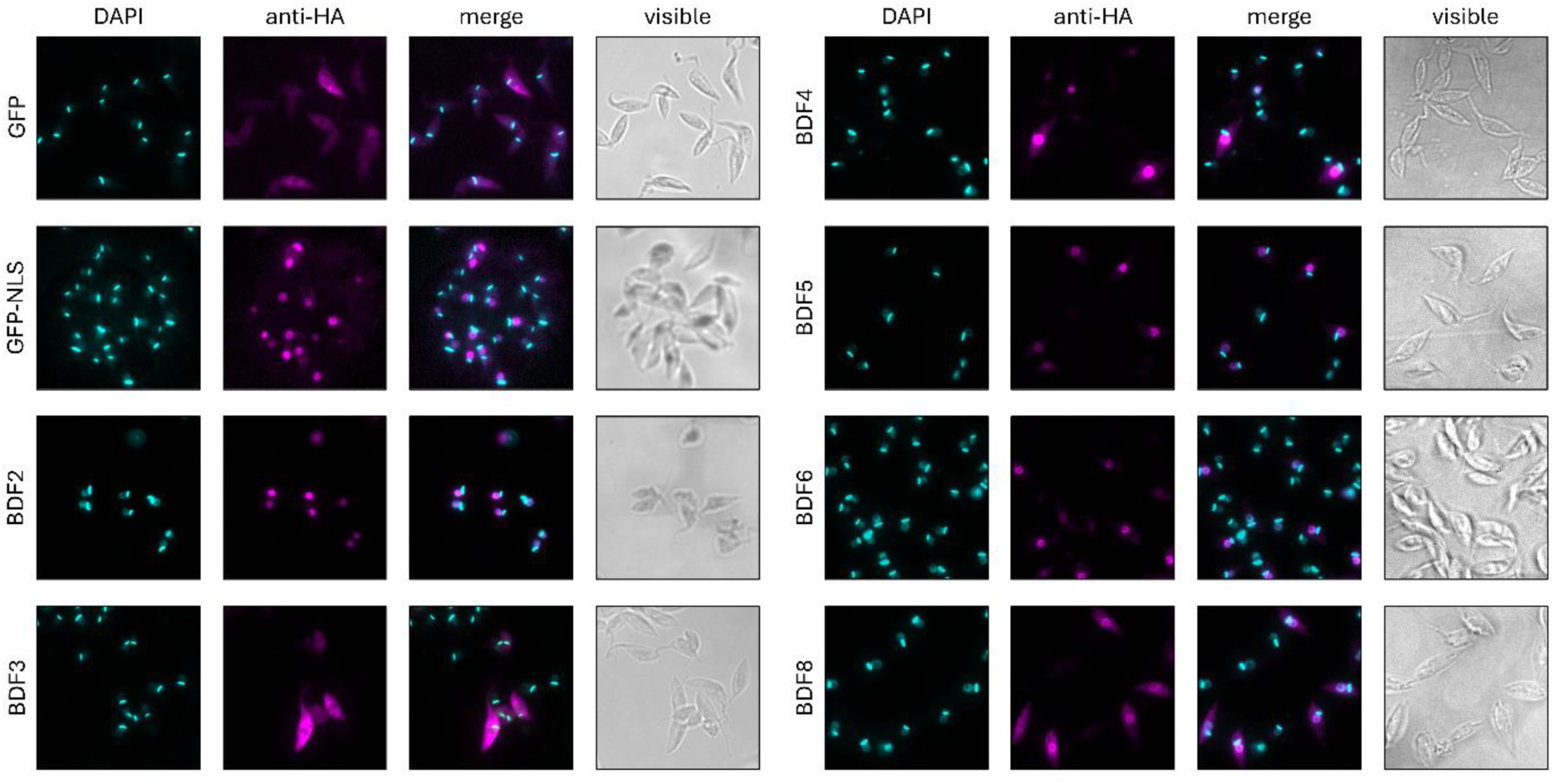
Subcellular localization of TurboID fusions determined by IFI. Columns show the DAPI signal in cyan, the anti-HA signal in magenta, the combined signal in both colors (merge), and the visible channel in grayscale. Each row correspond to a different linage. Expression conditions are those determined experimentally by western blot.

### Biotinylated proteins enrichment tests and mass spectrometry

We performed the purification tests using streptavidin-coated magnetic beads. For in vivo proximity biotinylation, we set the incubation time at 24 hours, ensuring coverage of all possible protein interactions during the different phases of the cell cycle. Parasites were incubated using the optimized expression conditions, crude extracts were prepared and the purification process was carried out, separating fractions at different points to evaluate the efficiency of the procedure.

Western blot analysis using streptavidin-HRP demonstrated the presence of biotinylated proteins in the inputs (IN), their absence in the flow-throughs (FT), and their successful enrichment in the eluates (EL), confirming the effectiveness of the purification process (**Supplementary Fig. 3A,B**). Only the TurboID-BDF2 linage showed signal in the FT, indicating saturation of the magnetic beads. The banding patterns of biotinylated proteins detected in the eluates were similar to those of their corresponding inputs, while their profiles differed slightly when comparing between linages. Silver staining of the eluates provided further evidence of biotinylated protein enrichment (**Supplementary Fig. 3C,D**). Samples from linages expressing TurboID fusions showed clear protein enrichment, while those from the negative control linage (without TurboID) were depleted in proteins. It is worth mentioning that the eluate of BDF2 showed the strongest intensity, followed by BDF8.

Subsequently, we scaled up the entire process to obtain sufficient sample for label-free quantification (LFQ) mass spectrometry analysis, processing each linage in quadruplicate. The linages were distributed across two experimental rounds, both containing the two spatial controls, as shown in **Supplementary Table 1**.

### BDF1 y BDF3 are detected outside the nuclear proximity proteome

Mass spectrometry results were processed to identify and quantify proteins present in the samples [34]. Resulting tables with normalized intensities from both experimental rounds (**Supplementary Files 1 and 2**) were processed and analyzed by hierarchical clustering and principal component analysis (PCA), followed by statistical tests to detect significant differences [35].

Heatmaps generated by hierarchical clustering of Z-score-normalized intensities show that all samples clustered by linage, indicating good correlation between replicates and experimental reproducibility (**Supplementary Fig. 4**). For the first experimental round, control samples lacking TurboID showed very low protein abundance (**Supplementary Fig. 4A**) and clustered far from the rest of the samples, which is expected. As their low values masked the visualization of differences between other replicates, we removed these samples and re-normalized the data to generate another heatmap (**Supplementary Fig. 4B**). All samples for BDF2, BDF5, BDF6 and BDF8 clustered in the same branch containing nuclear spatial control samples, while the cytoplasmic control samples were separate. For the second experimental round, BDF4 samples clustered with the nuclear spatial control, while BDF3 samples clustered with the cytoplasmic spatial control (**Supplementary Fig. 4C**). This is consistent with the subcellular localizations observed by fluorescence microscopy (**Fig. 1**) and previous reports [15]. Similarly, PCA shows that samples from BDF fusions that were detected in the nucleus by IFI cluster around the nuclear spatial control, while BDF3 samples are closer to those of the cytoplasmic control (**Supplementary Fig. 5**).

The experimental design also aimed to define the proximity-based nuclear proteome of *T. cruzi*. For this purpose, enrichments from the nuclear spatial control (GFP-NLS) were simultaneously compared against both the cytoplasmic spatial control (GFP) and the negative controls (NoTurbo) generated in the first experimental round (**Supplementary Fig. 6A**). To validate the method’s specificity, the distribution of proteins annotated with terms related to the nucleus was analyzed. When searching for proteins containing the words “nuclear”, “nucleus”, “nucleolar”, or “chromosome”, the vast majority were found to be overrepresented in the nuclear spatial control samples (**Supplementary Fig. 6B**), indicating that this control was indeed enriched with nuclear proteins. For this experimental round, proteins comprising the proximity-based nuclear proteome were identified as those significantly enriched (FDR=0.01) in GFP-NLS compared simultaneously to GFP and NoTurbo (**Supplementary Fig. 6C**). Regarding bromodomain factors, BDF2, BDF5, BDF6, and BDF7 were significantly enriched in this proteome. Although BDF8 was detected in the samples, it did not meet the established statistical significance criteria. It is important to note that BDF1, BDF3, and BDF4 were not detected in any of the samples analyzed in this round.

For the second round, we did not include a control without TurboID, so we only compared the significance of LFQ intensities between the nuclear and cytoplasmic spatial controls (**Supplementary Fig. 6D**). In this nuclear proteome (FDR=0.01), BDF2, BDF4, BDF5, BDF6, BDF7, and BDF8 were significantly enriched. Notably, although BDF1 and BDF3 were detected in these samples, they did not show significant nuclear enrichment, suggesting a predominantly cytoplasmic localization or a more intricate distribution between the two cellular compartments.

To unify results, we classified the entire *T. cruzi* proteome into nuclear confidence categories by integrating the statistical significance values from both experimental rounds (**Fig. 2**). Proteins were categorized according to their detection status and significance in each round and then combined to assign six confidence levels (**Fig. 2A**). For example, proteins detected with a significance of FDR ≤ 0.01 in both experiments were assigned to the “Very High” nuclear confidence group (dark green lines); or “High” if detected with FDR values ≤ 0.01 in one experiment and between 0.01 and 0.05 in the other (light green lines). In other words, these nuclear confidence levels reflect the consistency and magnitude of the statistical significance of the enrichment.

**Fig. 2:**
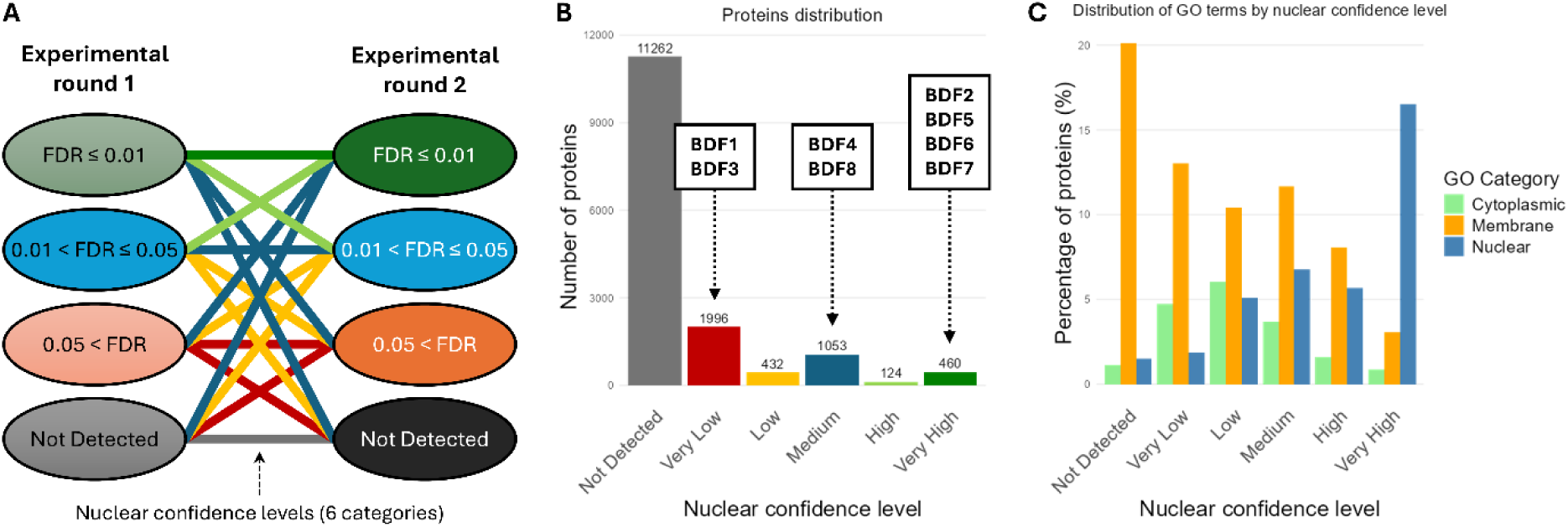
Distribution of proteins and GO annotations in the different nuclear confidence categories. (**A**) Schematic classification of the *T. cruzi* proteome based on the confidence levels of each experiment. The colored ellipses represent the proteins classified according to their detection and FDR values in each experimental round. The colors of the lines connecting them represent the confidence classification groups. (**B**) Distribution of the proteome in the different nuclear confidence categories. Colors for each category are the same as the lines in panel A. The number of proteins in each category is shown above each bar. (**C**) Percentage distribution of cytoplasmic (green), membrane (yellow), and nuclear (blue) GO terms in each nuclear confidence category.

The distribution of the proteome across these categories is shown in **Fig. 2B**, indicating the assigned category of each BDF. The analysis of the relative distribution of GO (Gene Ontology) annotations for cellular components validate the assignments of these categories (**Fig. 2C**) [36]. For example, the percentage of proteins with nuclear GO terms is minimal in the “Not Detected” category and gradually increases toward the “Very High” nuclear confidence category. Proteins with cytoplasmic GO terms are concentrated in the “Low” and “Very Low” confidence categories, being minimal in the “Very High” category. Membrane proteins are predominantly distributed in the low confidence categories, with a maximum in the “Not Detected” category. The proteome annotation according to these levels can be found in **Supplementary File 3**.

These results are consistent with the localization patterns of BDF fusions, indicating that TurboID does not alter their native distribution. For BDF2, BDF5, BDF6, and BDF7, the methodology indicates a very high probability of nuclear localization, while BDF1 and BDF3, consistently detected as extranuclear, are classified as having a very low nuclear probability (**Fig. 2B**). Notably, BDF4 and BDF8 may exhibit more complex localizations, as they are classified as having medium probability, which may suggest a dual nucleocytoplasmic localization.

### The tested BDFs form two distinct proximity networks with shared components

To build the proximity interactome of each BDF, we statistically compared the normalized intensities from the bait samples against both spatial controls (**Fig. 3**). Significantly enriched proteins against both controls (FDR_GFP_ < 0.05 and FDR_GFP-NLS_ < 0.05) were separated into two groups. One group, that we called “interactors”, contained those proteins that simultaneously showed high enrichment (T-test difference > 2) with respect to both spatial controls. The second group, that we called “proximals”, contained those proteins that showed high statistical significance against both controls (maximum q-value < 0.001), but only exceeded the enrichment threshold against one of the spatial controls or were close to the established threshold. In other words, we recovered those proteins that could be considered physical interactors given their high levels of enrichment and those that could be found in their proximity due to their statistical consistency. Detailed results can be found in **Supplementary File 4**.

**Fig. 3:**
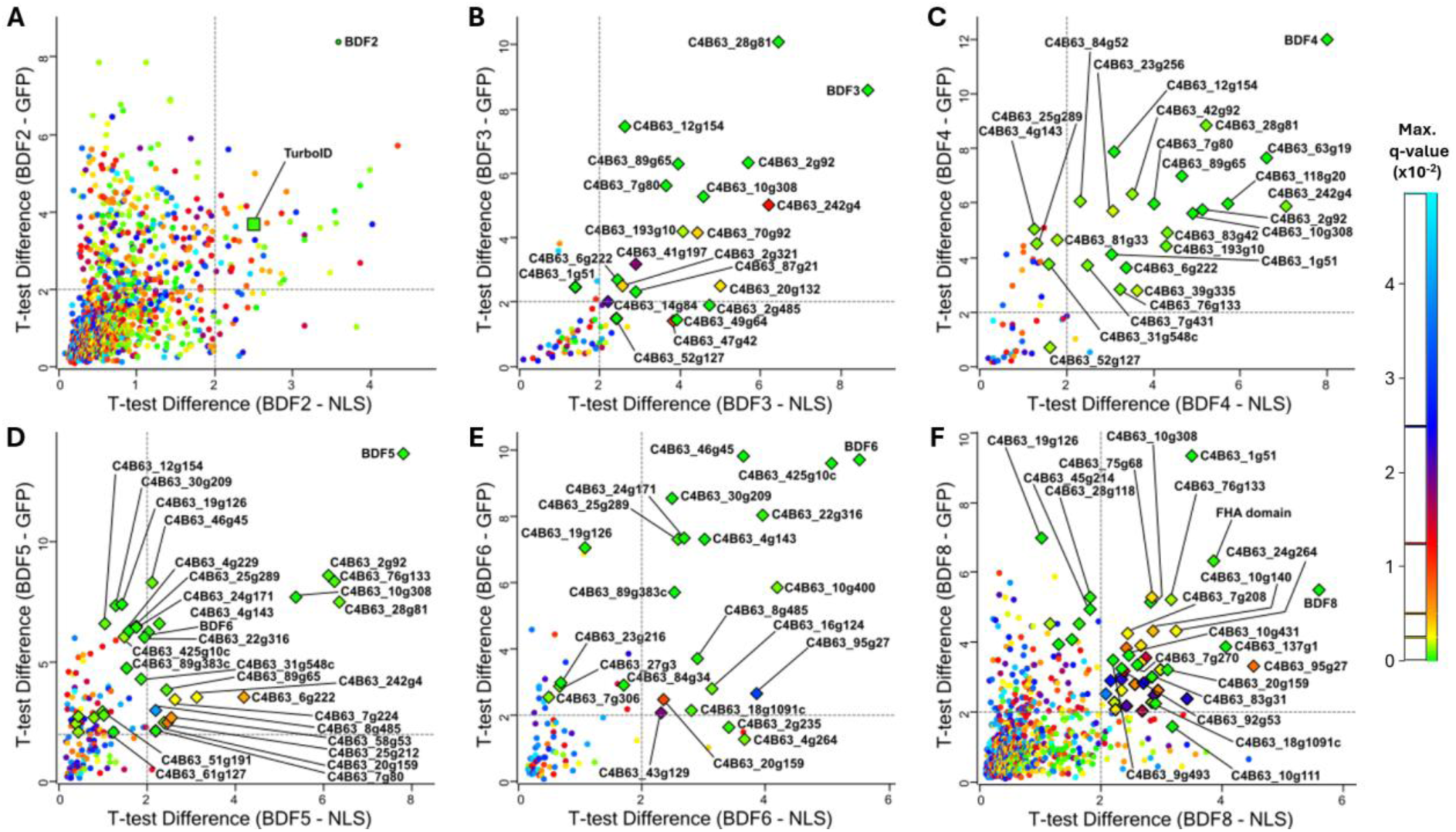
Proteins significantly enriched with respect to both spatial controls. Scatter plots comparing the signal difference of each protein between the bait and the cytoplasmic spatial control (BDF-GFP) versus the signal difference between the bait and the nuclear spatial control (BDF-NLS). The color scale represents the maximum q-value (FDR) between both T-tests for each protein. Green colors are more significant (lower q-value) and blue less significant (higher q-value). Proteins with maximum q-values > 0.05 were removed. Only differences greater than zero are shown. Proteins highlighted with diamonds were used for network analysis, and some of their identifiers are shown. Gray dotted lines indicate the enrichment cutoff values used to consider interactors. (**A**) Proteins significantly enriched in samples from parasites expressing the BDF2 fusion. It can be observed that the fusion gene was overexpressed compared to the controls, as TurboID was detected as a highly enriched protein. Significantly enriched proteins for BDF3 (**B**), BDF4 (**C**), BDF5 (**D**), BDF6 (**E**) and BDF8 (**F**).

The analysis revealed different levels of expression among the fusions. In the case of BDF2, TurboID was significantly enriched (**Fig. 3A**), suggesting that TurboID-BDF2 expression levels were excessively high and may have masked specific interactions. Therefore, BDF2 data was disregarded. BDF8 also showed a profile with numerous significantly enriched proteins (**Fig. 3F**). However, unlike BDF2, BDF8 did not exhibit differential enrichment of TurboID and maintained a specific interaction pattern, as it shared many proteins also enriched in BDF3, BDF4, BDF5, and BDF6 samples (see below). The remaining fusions showed more selective enrichment profiles (**Fig. 3B**-**E**).

Using these results (**Supplementary File 4**), we build an interaction graph representing interacting pairs with solid edges and proximal pairs with dashed edges, reflecting the measured enrichment with line thickness (**Fig. *4***). We represented the node degree with different colors, indicating the number of baits for which each protein was significantly enriched. Also, we added the nuclear confidence category (**Supplementary File 3**) as a colored bar. The network shows a highly interconnected group of proteins involving BDF3, BDF4, BDF5, and BDF8, associated with seven other proteins: HAT2, ENT, BDF5BP, 89g65, 242g4, 10g308, 76g133, and 10g308. In contrast, the connectivity pattern of BDF6 is different, being enriched primarily with exclusive proteins. Several BDF6 interactors can be identified as homologous components of the yeast NuA4 complex [37]. These include Yaf9/YEA2, EAF6, and HAT1, the latter being homologous to Esa1, the catalytic component of NuA4 [38]. TcTINTIN components MRGx and MRGBP are also classified as BDF6 interactors [21]. However, BDF6 is not completely isolated from the rest of the network, as it exhibits proximity to the other BDFs, particularly to BDF5 (classified as interactor), and some proximal proteins shared with BDF4 and interactors with BDF8. It is worth noting that no HDAC was found in these proximity proteomes.

**Fig. 4:**
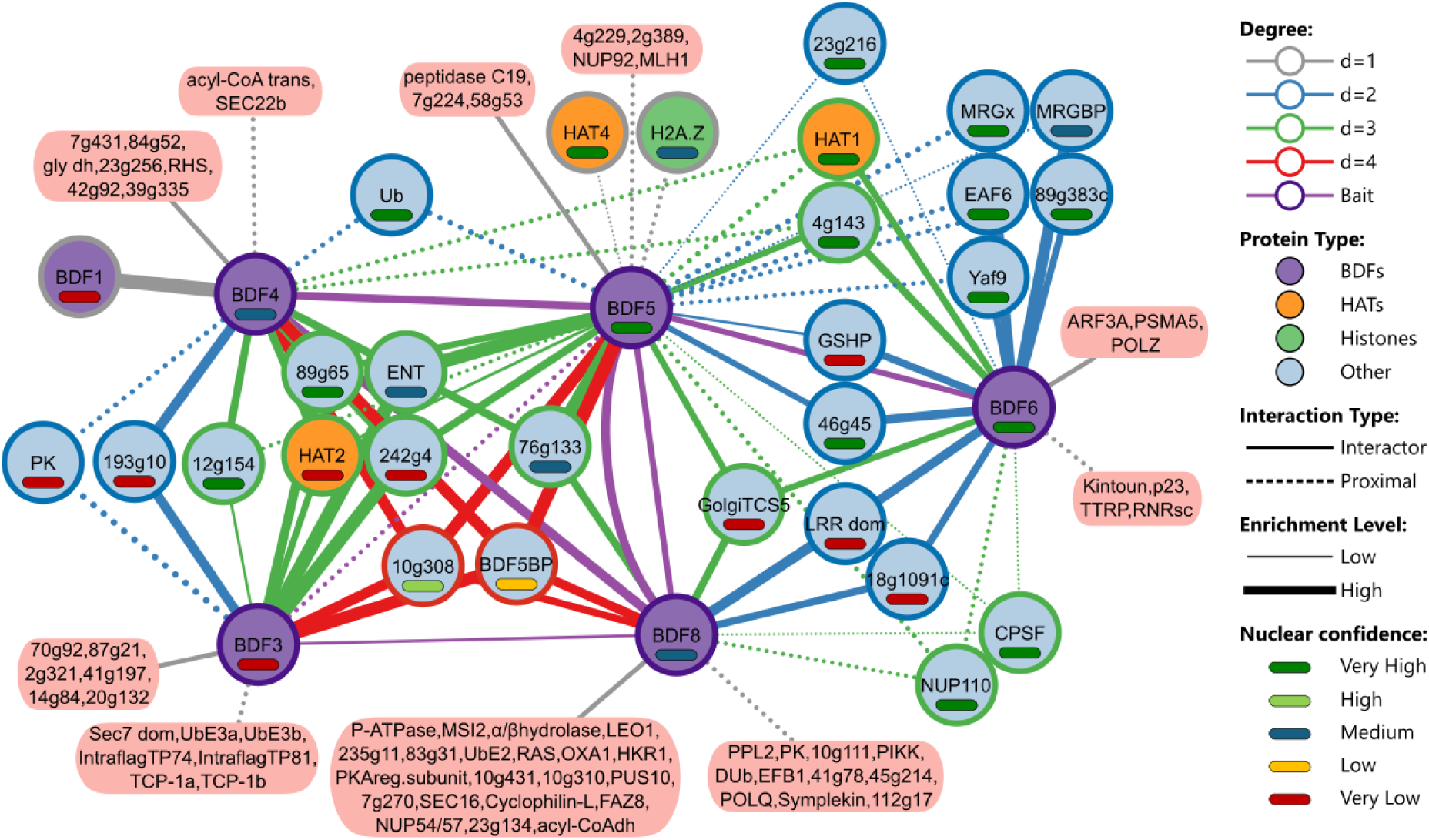
Proximity network of *T. cruzi* BDFs. The network was built using Cytoscape from the PL data. Nodes represent proteins, with BDFs shown in purple. Protein pairs classified as interactors are represented with solid lines and proximals by dashed lines. Line thickness reflects the magnitude of enrichment relative to the nuclear control. Line color indicates the degree of interconnectivity (i.e., the number of experiments in which the protein was detected). The nuclear confidence of each protein is indicated by a colored bar.

It can also be seen that BDF1 does not seem to be an integral part of the subnetwork that involves BDF3, BDF4, BDF5, and BDF8. Instead, BDF1 is highly enriched as an interactor of BDF4. This suggests that the interaction between BDF1 and BDF4 is not simultaneous with the formation of a larger complex involving the other components. Conversely, the connections established by BDF3 indicate that it must be an integral component. In contrast, the intraflagelar transport proteins TP74 and TP81 are detected near BDF3, consistent with observations that associate it with the cytoskeleton and the flagellum [15]. Notably, many components of this subnetwork have intermediate or very low nuclear confidence, including HAT2 (very low), in line with the subcellular localization of BDF3. We hypothesize that nascent histones are acetylated in the cytoplasm and subsequently imported into the nucleus, similar to H4K12 acetylation in yeasts [39]. If so, extranuclear BDFs could play a role in this process.

Among the interactors of BDF5 are two previously reported partners: BDF8, which is also reciprocally enriched, and BDF5BP [21]. Proximal proteins include HAT4 and the variant histone H2A.Z. It is possible that H2A.Z was enriched due to interactions between acetylated lysines and its BDs, since acetylated peptides of H2A.Z were recognized by *L. mexicana* BDF5 [6]. Shared components between BDF5 and BDF6 include HAT1, EAF6, Yaf9/YEA2, MRGx, MRGBP, GSHP, GolgiTCS5, 4g143, 89g383, and 46g45. These are highly enriched interactors of BDF6 and are classified as proximals to BDF5, except for GSHP, 4g143, and 46g45, that are classified as BDF5 interactors. All of them are high-confidence nuclear components, except for MRGBP (medium confidence), the conserved oligomeric Golgi complex subunit 5 (GolgiTCS5) and the glutathione peroxidase GSHP (very low). On the other hand, three extranuclear proteins are detected as shared interactors between BDF6 and BDF8: a protein with an LRR domain, the GolgiTCS5, and a hypothetical protein (18g091c).

The correspondence between the protein symbols used in **Fig. *4*** and gene identifiers can be found in **Supplementary File 5**.

Interactors of each BDF were also analyzed using Venn diagrams (**Supplementary Fig. 7** and **Supplementary Table 2**). Similarly to the interaction network, BDF3, BDF4, BDF5, and BDF8 share many interactors, while BDF6 share only some proteins with BDF5 and BDF8. Among all BDFs, BDF5 shows the greatest overlap in interactors, indicating that it is central hub protein.

### Domain architecture and predicted PPIs of T. cruzi CRKT and NuA4 complex components

The components we identified are orthologous to the identified components of the CRKT complexes in *L. mexicana* and *T. brucei*, and the NuA4 components identified in *T. brucei* [18–20]. While the stable components of the underlying complexes may vary depending on whether we consider proteins such as BDF1 as part of CRKT, **Table 1** and **Table 2** summarize those we considered to continue with the structural bioinformatics analysis.

**Table 1:**
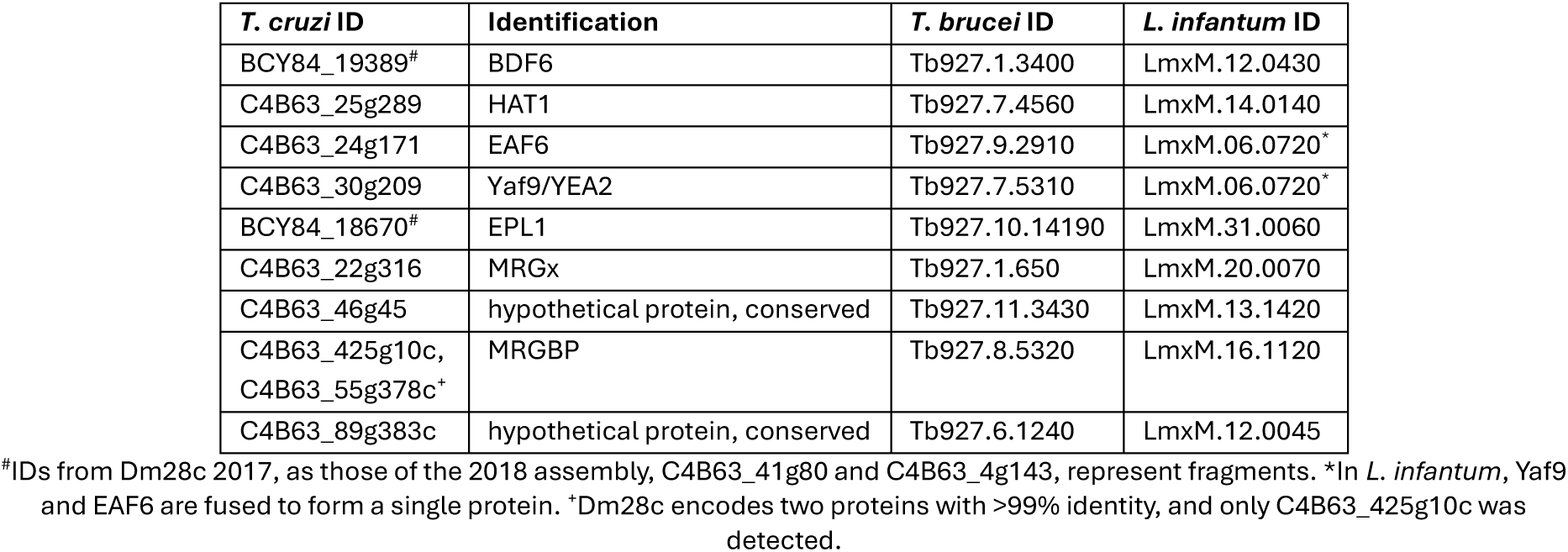
Components selected for the structural bioinformatics analysis of the subnetwork that would form the NuA4 complex. Genomic identifiers (IDs) of proteins detected by proximity labeling in *T. cruzi*, the identification made by sequence analysis or literature and the IDs of their orthologs in *T. brucei* and *L. mexicana*.

**Table 2:** Selected components for the structural bioinformatics analysis of the subnetwork that would form the CRKT complex. The genomic identifiers (IDs) of proteins detected by proximity labeling in *T. cruzi*, the identification made by sequence analysis or literature, and the IDs of their orthologs in *T. brucei* and *L. mexicana* are shown.

| *T. cruzi* ID | Identification | *T. brucei* ID | *L. mexicana* ID |
| --- | --- | --- | --- |
| C4B63_63g19 | BDF1 | Tb927.10.8150 | LmxM.36.6880 |
| C4B63_40g26 | BDF3 | Tb927.11.10070 | LmxM.36.3360 |
| C4B63_25g318 | BDF4 | Tb927.7.4380 | LmxM.14.0360 |
| C4B63_1g51 | BDF5 | Tb927.11.13400 | LmxM.09.1260 |
| C4B63_7g80 | BDF8 | Tb927.4.2340 | LmxM.33.2300 |
| C4B63_6g222 | HAT2 | Tb927.11.11530 | LmxM.28.2270 |
| C4B63_28g81 | ENT | Tb927.11.5230 | LmxM.24.0530 |
| C4B63_2g92 | BDF5BP | Tb927.9.13320 | LmxM.35.2500 |
| C4B63_242g4,<br>C4B63_76g133<sup>#</sup> | hypothetical protein, conserved | Tb927.7.2770 | LmxM.22.0880 |
| C4B63_89g65,<br>C4B63_202g8<sup>*</sup> | hypothetical protein, conserved | Tb927.6.1070 | LmxM.12.0230 |
| C4B63_10g308 | poly-ADP ribose polymerase domain (PARP) | Tb927.3.4140 | LmxM.08_29.1550 |

<sup>#</sup>Dm28c strain encodes two proteins with >99% sequence identity. Both were detected in proximity interactomes and we continued working with 242g4. <sup>\*</sup>This protein is also encoded by two genes. They have >99.5% identity, and only 89g65 was detected.

We processed the amino acid sequences of these proteins using MultimerMapper, developed in our laboratory for the systematic and automated analysis of multimeric structures predicted with AlphaFold (AF) [23–25]. Briefly, this iterative method begins by predicting all possible dimeric combinations. MultimerMapper analyzes these dimers and their confidence metrics (PAE, pLDDT, etc.) to identify interacting pairs. These interactions are then used as seeds to explore the stoichiometric space by suggesting new combinations for prediction. Successive iterations expand to trimers, tetramers, pentamers, etc., with an algorithm deciding which paths to explore by identifying combinations that produce cohesive complexes with no detached subunits. Each of these stable stoichiometries are capable of generating children stoichiometries by adding new subunits, and those where all of their offspring is unstable are considered convergent stoichiometries, since they no longer possess free interaction surfaces to assemble larger complexes.

In parallel, the program tracks stoichiometric context, PPI frequencies and conformational variability. This enables the detection of interaction and conformational changes associated with the presence or absence of certain proteins. We prioritized PPIs with the highest observed frequencies, because they were repeatedly observed across multiple modeling contexts, therefore making them the most likely true positives. We considered dynamic PPIs only when they provided mechanistic insights relevant to complex assembly or function. Finally, we examined the output stoichiometric space to derive the predicted stoichiometric coefficients for each complex.

After running MultimerMapper, we extracted the best local quality model of each protein (highest average pLDDT) and analyzed their corresponding intramolecular PAE array to detect segments with high internal cohesion, indicative of discrete globular domains. We next queried structural databases using the Foldseek structural similarity search algorithm [40] with their atomic coordinates to identify conserved domains and assign appropriate names to hypothetical proteins. Resulting domain architectures are shown in **Fig. 5A**-**B**. Three of the four unnamed proteins (**Table 1** and **Table 2**) yielded similar structures that could be considered homologs. The high-frequency (≥70%) PPI network predicted by MultimerMapper for NuA4 is shown in **Fig. 5C** and the full network for NuA4 and CRKT are presented in **Supplementary Fig. 8** and **Fig. 5D**, respectively.

**Fig. 5:**
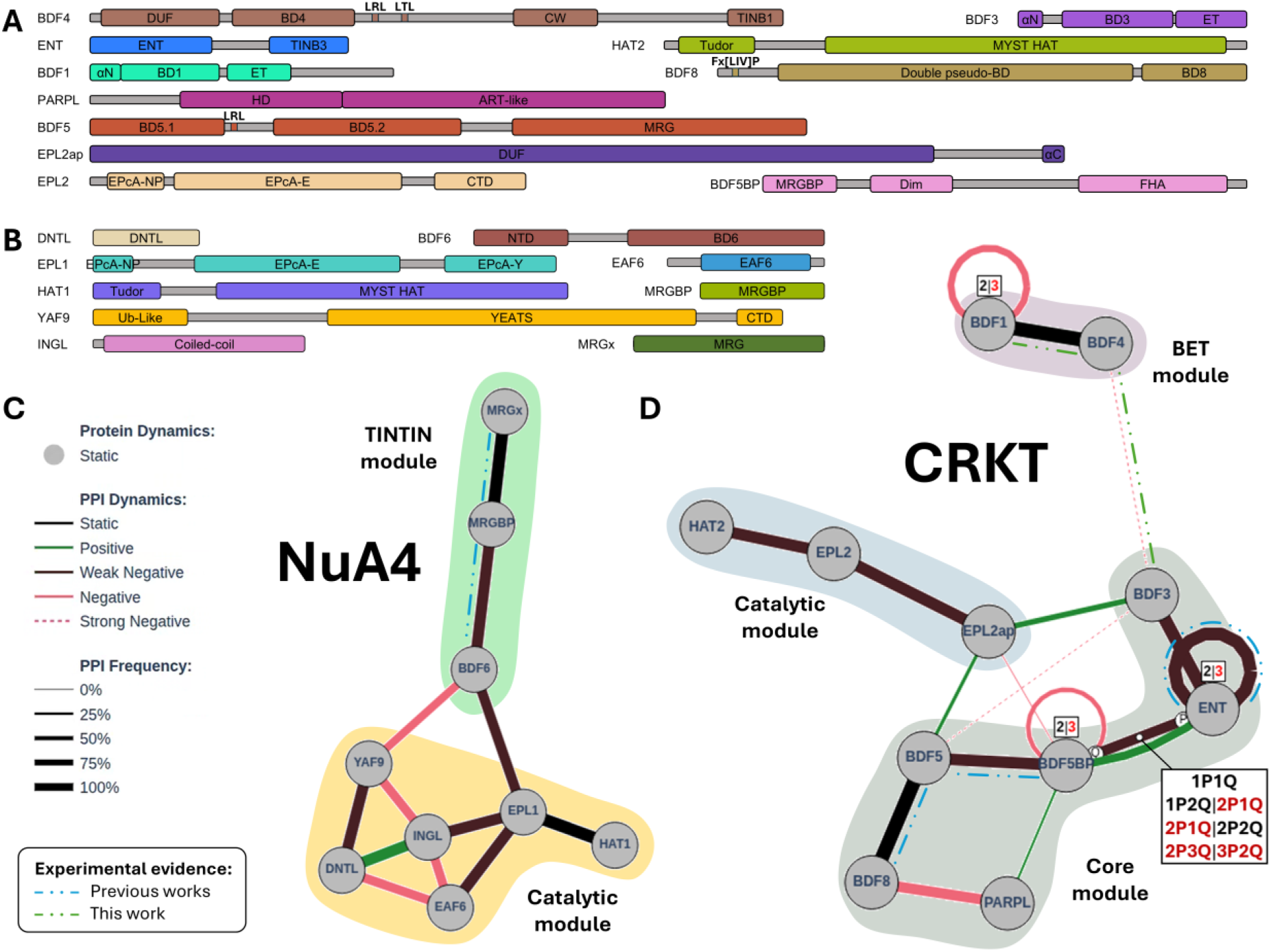
Identified domain architectures and predicted PPI networks using AlphaFold3 and MultimerMapper. Domain architectures identified with FoldSeek for proteins of the CRKT (**A**) and NuA4 (**B**) complexes. Predicted PPI graphs obtained with MultimerMapper for the protein subsystems of the NuA4 (**C**) and CRKT (**D**) complexes. Edges represent distinct PPI modes. Edge width is proportional to the frequency of detection of the PPI among all the predictions that contain the target pair. NuA4 PPIs were filtered to show only those with a frequency of at least 70%. Edge colors indicate if the PPI was detected in dimeric predictions (2-mers) for each protein pair (**Static**: always detected; **Positive**: not detected in 2-mer; **Negative**: detected in 2-mer) and the strength indicates the frequency (**Weak**: >80%, **Strong**: 0%) in more complex protein combinations (N-mers). Identified structural modules are indicated in different colors (see below). Boxes indicate stable (black) and unstable (red) combinations using the nPmQ nomenclature for multivalent interactions (more than one interaction site) and numbers for homooligomers. Direct interactions with experimental evidence are indicated with dashed lines.

Regarding the hypothetical components of NuA4 (**Table 1**), the best match for 46g45 was the model AF-B3H615-F1-model_v4 from *A. thaliana* (**Supplementary Fig. 9**). This protein is annotated as “PHD finger protein ING2”, belonging to the group known as Inhibition of Growth (ING) proteins, which have a coiled-coil segment of two antiparallel α-helices and an N-terminal PHD domain. These are part of histone acetylation-related complexes (HARCs) in humans, yeast, and plants, including the NuA4 complex [41]. Given the high structural similarity but the absence of the PHD domain, we named this protein INGL (ING-like). For 89g383c, we found the entire protein to be similar to the N-terminal domain of DMAP1 (DNA Methyltransferase 1 Associated Protein 1). This protein is involved in various HARCs and interacts with the C-terminal α-helix domain of YAF9 within the human SCRAP-C complex [42]. Since *T. cruzi* YAF9/YEA2 interacts with 89g383c via a high-frequency PPI (**Fig. 5C**) and only conserves homology with the N-terminal segment of DMAP1, we named this protein DNTL (DMAP1 N-terminal-Like).

For the hypothetical components of CRKT (**Table 2**), only 242g4 recovered a similar structure linked to HARCs. Its best hit was the model AF-Q9FX82-F1-model_v4 of the Enhancer of Polycomb-like (EPL) protein from *Arabidopsis thaliana*. EPL proteins are involved in the formation of complexes with HAT activity and interact directly with HAT enzymes through their EPcA-E domains [43], very similar to one of the globular segments of 242g4 (**Supplementary Fig. 9**). Furthermore, MultimerMapper predicts that it interacts with HAT2 through a high-frequency PPI (**Fig. 5D**). 242g4 conserves the first two EPcA subdomains (NP and E) of eukaryotic EPL proteins. EPcA-NP (<u>N</u>ucleosome core <u>p</u>article) is involved in the interaction with nucleosomes, and EPcA-E (<u>E</u>sa1) with HAT proteins. The third subdomain, EPcA-Y (Yng2/Eaf6), which in other EPLs is an α-helix that interacts with the Yng2 and Eaf6 components of the NuA4 complexes [43], is replaced in this case by an unidentifiable C-terminal domain (CTD). As EPL1 was already identified as a component of *T. cruzi* NuA4 (**Table 1**) [17], we named it EPL2. On the other hand, 89g65 recovered partial hits from eukaryotic importins, some involved in the import of histones from the cytoplasm to the nucleus [44], but their significance was unclear. Therefore, we decided to name it EPL2ap (EPL2-associated protein), according to the physical association with EPL2 predicted by MultimerMapper (**Fig. 5D**). EPL2ap is composed of a globular domain of unknown function (DUF) and a disordered segment that ends in a C-terminal α-helix (αC).

BDF1 and BDF3 are paralogous to each other (**Supplementary Fig. 10A**-**C**), each possessing an N-terminal α-helix (αN), a central BD and a C-terminal ET domain [13], where BDF1 has an extra disordered C-terminal segment. BDF4 has four globular segments: an N-terminal domain of unknown function (DUF) fused to its BD, followed by a CW domain [45] and a coiled-coil C-terminal domain (**Supplementary Fig. 10D**). ENT has been structurally characterized in *T. brucei*, confirming in vitro the dimerization capacity of its ENT domain (<u>E</u>MSY <u>N</u>-terminal). Also, a second coiled-coil C-terminal domain potentially involved in PPI was suggested [46]. The predicted structure of both domains can be seen in **Supplementary Fig. 10E**. Consistently, MultimerMapper predicts that ENT is capable to homodimerize, but not to homotrimerize (indicated by the inset over the self-interaction edge of ENT in **Fig. *5*D**). Meanwhile, two highly similar interactions modes are predicted for the BDF1-BDF4 and BDF3-ENT pairs, where the αN domains of the former engage the coiled-coil domain of the latter, respectively (**Fig. *5*D and Supplementary Fig. 10F-G**).

Since we had the coding sequences for BDF1 and BDF4 available in the laboratory, we decided to test the interaction of this pair and verify that it occurred through the C-terminal domain of BDF4. Yeast two-hybrid (Y2H) interaction assays demonstrated that the BDF1-BDF4 pair is capable of interacting using their complete sequences, but the interaction is disrupted upon removal of the αN domain from BDF1 (**Supplementary Fig. 11**), consistent with the interaction model predicted by MultimerMapper. Based on these observations, we decided to name the CTD domains of BDF4 and ENT as TINB1 and TINB3, for <u>T</u>rypanosomatid Interactor of <u>N</u>-terminal <u>B</u>ET 1 (BDF1) and 3 (BDF3), respectively (**Supplementary Fig. 10H** and **Fig. *5*A**).

The domain architecture and interactions between BDF5, BDF5BP, and BDF8 have been previously described [21] and their PPIs are properly identified by MultimerMapper (**Fig. *5*D**). BDF5 has two tandem N-terminal BDs separated by a disordered segment of a C-terminal MRG domain. BDF5BP has a MRG binding domain (MRGBP) that bind the MRG of BDF5, a central domain that MultimerMapper predicts to be involved in its homodimerization (Dim) and several dynamic interactions (see below), and a C-terminal FHA domain with the potential to recognize phosphorylated tyrosines/threonines [47]. BDF8 has a disordered N-terminal domain with a conserved Fx[LIV]P motif that interacts with the MRG domain of BDF5, a central DUF formed by two pseudo-BDs, and a C-terminal BD that partially conserves the acetylated lysine recognition pocket [21]. HAT2 has an N-terminal Tudor domain and a MYST HAT domain. In *T. brucei*, the Tudor domain is unable to recognize methylated lysines [48], while the MYST HAT domain acetylates histone H4 at K2, K5, and K10 [49]. Although the presence of a poly-ADP-ribose polymerase (PARP) domain has been identified with high probability in the orthologs of 10g308 in *T. brucei* and *Leishmania* [19, 50], Foldseek captures just partial structural similarity with this family. The *T. cruzi* genome encodes one PARP protein (**Supplementary Fig. 12A**) that highly similar to the PARP proteins of other eukaryotes (**Supplementary Fig. 12B**) and performs the same function of polymerizing ADP-ribose in response to DNA damage [51]. 10g308, on the other hand, conserves only the HD (<u>H</u>elical <u>S</u>ubdomain), but lacks the WGR (Tryptophan-Glycine-Arginine) domain. Meanwhile, the catalytic ART (<u>A</u>DP-ribosyltransferase) domain retains only partial structural characteristics, which is why we call it ART-like domain (**Supplementary Fig. 12C**). HD domains regulate the activity of ART domains through conformational changes that control NAD+ entrance to the catalytic site [52]. We think that 10g308 is likely a paralog of PARP that maintains some kind of HD domain functionality, while the ART-like domain has a unique trypanosomatid function, since it does not conserve the catalytic pocket and we have not found this type of fold in other organisms. For this reason, we decided to name this protein as PARPL (PARP-Like).

Regarding the domains of NuA4 components (**Fig. 5B**), unlike EPL2, EPL1 retains the three EPcA subdomains, but not the C-terminal segment that recognizes the TRA module in yeast [43]. Both EAF6 and HAT1 (Esa1 homolog) have the characteristic folds of their yeast homologs [53]. MRGx, MRGBP, and BDF6 have been previously described: MRGx and MRGBP interact with each other and consist of a single MRG domain and a single MRGBP domain, respectively; while BDF6 possesses a C-terminal BD that binds to MRGBP and an N-terminal domain [21]. The YEATS and CTD domains of YAF9/YEA2 are highly conserved [54], but an extra N-terminal domain with a ubiquitin-like fold (Ub-like) is present, which could play a role in regulating proteasome activity, a phenomenon observed in other proteins with Ub-like folds [55]. This domain is also highly similar to the N-terminal domain of human coilin, a nuclear protein that is part of Cajal bodies and whose N-terminal domain has an Ub-like fold essential for its self-assembly and the formation of these nuclear bodies, non-membrane organelles involved in the assembly and maturation of ribonucleoprotein complexes [56]. This suggests that the Ub-like domain of YAF9 could also mediate specific protein-protein interactions within NuA4, analogous to the function of the coilin N-terminal domain in Cajal body assembly.

The high-frequency PPI network for NuA4 components (**Fig. 5C**) captures both PPIs involved in TcTINTIN formation: MRGx-MRGBP and MRGBP-BDF6 [21]. Meanwhile the remaining components form a denser PPI network that we identified as the catalytic module of NuA4 (see below). Notably, TcTINTIN binds to this module through a PPI between BDF6 and EPL1, consistent with previous NuA4 descriptions in yeast and humans [22].

On the other hand, the PPI network for CRKT components (**Fig. 5D**) is divided into three distinct high-frequency PPI subnetworks, connected through more dynamic, low-frequency interactions. One subnetwork corresponds to a catalytic module (see below) and involves HAT2, EPL2, and EPL2ap. Another subnetwork that we designated as the core module of CRKT (see below) includes BDF3, ENT, BDF5BP, BDF5, BDF8, and PARPL. Within this subgroup, besides the dimerization of ENT, the PPIs between the pairs BDF5-BDF5BP and BDF5-BDF8 are also detected, as previously reported [21, 46]. ENT and BDF5BP establish multivalent interactions (more than one interaction surface) that allow the assembly of a heterotetrameric subcomplex with a 2:2 stoichiometry. The last subnetwork contains the interacting pair BDF4-BDF1 and connects through dynamic interactions between BDF4 and BDF3. Since this module involves interactions between the two members of the *T. cruzi* BET family, BDF1 and BDF3, we designated it as BET module.

### Stoichiometry and structural inference predicts a piccolo-like NuA4 and a unique CRKT complex organized around a central symmetry-core

The stoichiometric space exploration algorithm of NuA4 sequences (**Table 1**) converges on a complex with a 1:1:1:1:1:1:1:1:1 stoichiometry (**Fig. *6*A** and **Supplementary Fig. 13**). In this complex, MRGx, MRGBP, and BDF6 form the TcTINTIN module, where BDF6 extends its CTD domain into the catalytic module via a flexible region, engaging with the C-terminal portion of the EPcA-Y domain of EPL1. This type of interaction has already been observed between EPL1 and BDF6 homologs [57]. Simultaneously, the CTD domain of BDF6 interacts with the Ub-Like domain of YAF9/YEA2. Considering that the Ub-Like domain participates in a PPI within the complex suggests a similar function to that of the N-terminal domain of coilin, involved in PPIs essential for Cajal body assembly [56], rather than to proteasome modulatory functions [55].

**Fig. 6:**
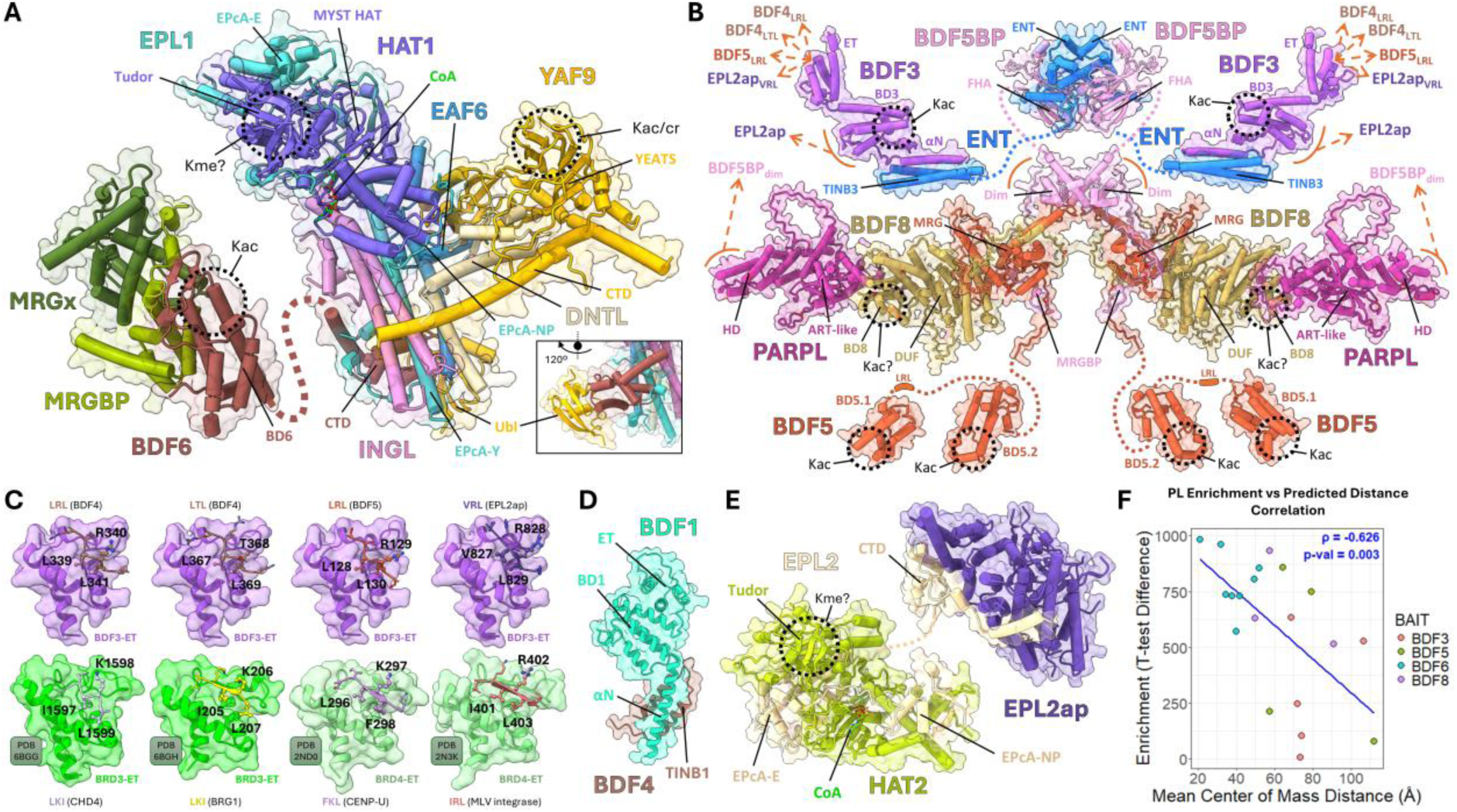
Predicted structures of NuA4 and CRKT complexes correlates with proximity data. The protein names and their domains are indicated using the same colors schemes as in Fig. 5. Dotted circles indicate potential recognition pockets for post-translational modifications. Most of the disordered loops were removed to improve visualization. (**A**) NuA4 AF3 model of the convergent stoichiometry detected by MultimerMapper. The cofactor molecule Coenzyme A (CoA) was transferred from the PDB model 1MJA to HAT1. The inset shows a side view of the interaction between the CTD domain of BDF6, EPL1, and YAF9. (**B**) Combinatorial assembly of the CRKT core module using CombFold and stoichiometry 2:2:2:2:2:2. Dynamic interactions are indicated with dashed orange arrows. (**C**) MultimerMapper-predicted interaction modes between the ET domain of BDF3 (purple) and the LRL and LTL motifs of BDF4 (brown), the LRL motif of BDF5 (orange) and the VRL motif of EPL2ap (violet), compared with the interaction modes of the experimental structures of the ET domain of the human proteins BRD3 (green) and BRD4 (light green). PDB identifiers and residues involved in the interactions are indicated. (**D**) MultimerMapper-predicted interaction mode between the αN domain of BDF1 (light blue) and the TINB1 domain of BDF4 (brown). (**E**) AF3 model of the CRKT catalytic module, containing HAT2, EPL2, and EPL2ap, with the 1:1:1 convergent stoichiometry predicted by MultimerMapper. The CoA molecule was transferred from 1MJA to HAT2. (**F**) Correlation graph between protein enrichment levels in PL experiments and the average distances between the protein centers of mass computed in the proposed structural models. Colors correspond to the bait from which the enrichment level was taken. The value of the Spearman correlation coefficient (ρ), its statistical significance (p-value), and the fit of the linear model (blue line) are indicated.

The predicted catalytic (or HAT) module of NuA4 exhibits an architecture that integrates multiple subunits organized hierarchically (**Fig. *6*A**). INGL, the EPcA-Y domain of EPL1, and EAF6 are organized into four tightly packed α-helices that form the structural scaffold of the module, to which the remaining components are anchored. The catalytic component, HAT1, is anchored to the opposite end to which the CTD domain of BDF6 is attached, interacting tightly to the EPcA-E and EPcA-NP domains of EPL1, stabilizing it over the tip of INGL and EAF6. We located the catalytic site of HAT1 by transferring the Coenzyme A (CoA) from the crystal structure of its yeast homolog Esa1 [58] onto our model. The resulting alignment positioned CoA above a solvent-exposed surface adjacent to the Tudor domain. YAF9/YEA2 stabilizes its C-terminal α-helix domain through interactions with DNTL, which forms a bridge that anchors the YEATS domain to the EPL1-EAF6-INGL helical scaffold. The spatial distribution of these proteins is highly similar to the organization of piccolo-NuA4 from *S. cerevisiae* (a NuA4 subcomplex where TINTIN and TRA modules are absent), particularly in the disposition of its four α-helix scaffold (**Supplementary Fig. 14**) [59]. In *T. cruzi*, INGL fulfills a structural role similar to that of yeast YNG2. The only significant differences between these two complexes are the relative orientation of their catalytic subunits and the absence of YAF9/YEA2 and DNTL in piccolo-NuA4 from yeasts.

It is important to mention that we detected other three convergent stoichiometries (**Supplementary Fig. 13**). Two of them are smaller versions of the complex reached through different assembly paths, like the catalytic module without TINTIN. The third involves two subunits of EAF6 and one of each BDF6, EPL1 and HAT1. This last alternative involve PPIs detected with much less frequency, like the homodimerization of EAF6 (**Supplementary Fig. 8**). In contrast, the model in **Fig. *6*A** involves PPIs that were consistently detected and is the biggest convergent stoichiometry we found. Moreover, its high similarity to piccolo-NuA4 indicates that it is the most plausible representation of the complex operating in physiological contexts.

For CRKT, we were unable to reach convergent stoichiometries across all branches of the stoichiometric space because some identified complexes were too large and exceeded the AF3 server’s token limit (approximately 5000 amino acids) [60]. Instead, the model from **Fig. *6*B** is the result of the combinatorial assembly using CombFold [61] with core module proteins (**Fig. *5*D**) and the 2:2:2:2:2:2 stoichiometry. We propose this stoichiometry based on the architecture of the central heterotetramer formed by ENT and BDF5BP that has a 2:2 ratio (**Supplementary Fig. 15**). This subcomplex has two binding sites to engage with two subunits of BDF5 through the MRGBP domain of BDF5BP and two binding sites to engage two subunits of BDF3 through the TINB3 domains of ENT (**Supplementary Fig. 15A**). Since the remaining components from the core module establish high-frequency PPIs in a 1:1 ratio, the overall stoichiometry where all these interaction surfaces are covered results in 2:2:2:2:2:2. The resulting assembly exhibits central symmetry. BDF5BP, in addition to its indirect dimerization through interactions with ENT, homodimerizes via its central domain (Dim), whose center contains symmetry axis. From this central structure and outwards, BDF5 engages the MRGBP domain of BDF5BP and, to a lesser extent, with Dim. BDF8 also binds to the MRG domain via its bipartite interaction [21], which in turn binds to PARPL through interactions with two α helices of BD8. This leaves the potential acetyl-lysine recognition pocket of BD8 exposed to the solvent. Regarding the structure formed between ENT and BDF5BP, two ENT domains engage with two FHA domains, forming a globular architecture that exposes two phosphorylated residue recognition sites to the solvent (**Fig. *6*B** and **Supplementary Fig. 15B**), likely involved in signaling pathways. This structure is connected to the rest of the complex via disordered segments of BDF5BP, while the TINB3 domain of the ENT protein binds to BDF3.

While less confident due to their PPI frequencies, MultimerMapper predicts that the ET domain of BDF3 would be able to recognize linear motifs formed by two leucines or one valine and one leucine flanking either a threonine or an arginine ([LV][TR]L) in BDF4, BDF5 and EPL2ap (**Fig. *6*C**). Comparing these binding modes with those of human ET domains, which also recognize motifs formed by two hydrophobic amino acids flanking a hydrophilic amino acid [62], reveals the same conformation (**Fig. *6*C**). The flanking hydrophobic residues are in contact with the cavity of their respective ET domains and the central hydrophilic residue is exposed to the solvent. Y2H assays validate the existence of a weak interaction between BDF3 and BDF4 (**Supplementary Fig. 11**), which would connect BDF4 to the core module. Meanwhile, BDF4 binds BDF1 through its TINB1 domain (**Fig. *6*D**), forming the BET module. However, this engagement would be non-simultaneous to the interaction with the core module (see above).

The catalytic module of CRKT converges in a complex with 1:1:1 stoichiometry (**Fig. *6*E**), were EPL2 acts as a bridge between HAT2 and EPL2ap. The interaction between EPL2 and HAT2 conserves the binding mode of EPL proteins, where the EPcA-NP and EPcA-E domains wraps around HAT2. This is similar to how EPL1 binds to HAT1 in our *T. cruzi* models and how EPL1 interacts with Esa1 in yeasts (**Supplementary Fig. 16**). In the case of EPL2, its CTD domain replaces the typical EPcA-Y domain of EPL proteins and is responsible for binding EPL2ap. While predicted with less confidence, this module binds to the core through dynamic interactions between EPL2ap and BDF5, BDF5BP and BDF3 (**Fig. *5*D**). The interaction with BDF5BP is mediated through the C-terminal α-helix of EPL2ap and is the only one detected in simple dimeric predictions (**Supplementary Fig. 17A,B**). This interaction is predicted to activate the engagement of BDF5 (**Supplementary Fig. 17C**), but it is not detected when two subunits of BDF5BP are present, suggesting that this interaction is not possible when the Dim domain of BDF5BP is dimerizing. Subsequently, this interaction would not be present in the proposed model from **Fig. *6*E**. Nonetheless, after the DUF domain of EPL2ap, we found aminoacid sequence conservation only in the region of the α-helix that mediates these PPIs (**Supplementary Fig. 17D**), which supports the dynamic interaction model. Alternatively, besides interacting with the ET domain of BDF3, EPL2ap is predicted to engage its BD though the opposite surface of the acetyl-lysine recognition pocket (**Supplementary Fig. 18**). Both interactions could explain the enrichment of the catalytic module in BDF3, BDF5 and BDF8 samples (**Fig. *4***).

Another less confident interaction is the engagement between PARPL and the Dim domain of BDF5BP (**Fig. *5*D**). This interaction only appears when two subunits of BDF5BP are present in the predictions with at least one subunit of PARPL and is mediated mainly through the ART-like domain of PARPL (**Supplementary Fig. 19A,B**). Extra AF3 predictions using two subunits of each BDF5, BDF5BP and PARPL indicate that two PARPL subunits can coexist bound to Dim, while the tip of the MRG domain of BDF5 interacts through a different surface of Dim (**Supplementary Fig. 19C**). Similarly, we found that BDF1 is predicted to homodimerize in 50% of the models (**Fig. *5*D**).

Several studies have proposed that there is a labeling radius surrounding the baits in proximity labeling methodologies, defining the distance limit for biotinylation, implying that nearby proteins will be biotinylated to a greater extent than those farther away [63, 64]. Based on this idea, we statistically tested whether the distances between proteins in our structural models (measured as the separation between their centers of mass) correlate with their enrichment levels relative to the baits (**Fig. *6*F**). Spearman’s rank correlation analysis indicates a significant correlation (p-value = 0.003) of moderate to strong magnitude (ρ = −0.623). This indicates that proteins that we found more enriched for a particular bait are, in turn, closer in space within our structural models with respect to those that are less enriched. This observation gives statistical support to our methodology.

Finally, given the observed spatial proximity between NuA4 and CRKT (**Fig. *4***), we decided to systematically analyze the cross-interactions between the components of both complexes. Pairwise predictions revealed a single link connecting both complexes: a predicted PPI between EPL1 and BDF5 (**Supplementary Fig. 20A**). This observation is consistent with the fact that EPL1 is the most enriched NuA4 component in the BDF5 proximity proteome (**Fig. *4***). The interaction involves an unstructured region of EPL1 on the opposite face to the catalytic site of HAT1 within our NuA4 model (**Supplementary Fig. 20B**). This segment is close to the EPcA-E domain of EPL1 and engages an unstructured segment near the MRG domain of BDF5, forming an antiparallel β-sheet (**Supplementary Fig. 20C**). Since this segment is involved in two distinct interactions (with HAT1 and with BDF5), we explored the dynamic behavior of the BDF5/ELP1/HAT1 system to assess whether they are capable of coexisting (**Supplementary Fig. 20D**). The results indicate that the EPL1-BDF5 interaction becomes destabilized when HAT1 is introduced into the system. That is, according to the predictions of AlphaFold3 and MultimerMapper, EPL1 detaches from BDF5 in the presence of HAT1, preferentially binding to the latter.

## Discussion

In this work, we analyzed the PPIs mediated by five of the eight *T. cruzi* BDFs and inferred the stoichiometry and structure of their associated complexes using a combination of wet- and dry-lab approaches. Through the generation and analysis of PL data, we found that BDF3, BDF4, BDF5, and BDF8 form a dense proximity network with 6 other proteins. These components are orthologs to those identified by co-IP of *T. brucei* and *L. mexicana* BDFs [18–20], evidencing that the underlying CRKT complex is conserved in *T. cruzi*. We described yet another complex formed by 9 proteins organized in two modules. Its catalytic or HAT module is structurally similar to the piccolo version of NuA4 from yeasts, while the second module is TcTINTIN, which engages the HAT module though the CTD of BDF6.

Studies in *L. mexicana* indicate that CRKT is essential for RNA polymerase II-dependent transcription and parasite viability [20]. Using structural similarity searches, we assigned several CRKT components to conserved protein families, allowing us to infer potential molecular roles within the complex to proteins previously annotated as hypothetical. Based on similarities to EPL family members, we named one of these proteins as EPL2, suggesting that it may regulate HAT activity. Another component was identified as a PARP paralog and named it PARPL. This protein may possess unique trypanosomatid functions, as it is highly conserved among trypanosomatids but diverges from typical PARPs. However, we were unable to assign EPL2ap to any known protein family, suggesting that it represents a trypanosomatid-specific factor with unique functions within the complex.

Our proximity network indicates that the BDF1–BDF4 interaction is independent of CRKT, consistent with Co-IP data of *T. brucei* BDF1 and BDF4 [19], suggesting that this is a conserved feature of the complex. Using Y2H, we validated that this interaction occurs between the C-terminal domain of BDF4 and the αN domain of BDF1. A homologous interaction was also observed between the αN domain of BDF3, a BDF1 paralog, and the C-terminal domain of ENT. Based on these findings, we named the domains that engage BET family αN domains as TINB. These results provide a new framework to reinterpret previous reports of non-nuclear BDFs in *T. cruzi*. BDF1 and BDF3 were found out of the nucleus in *T. cruzi* [14, 15] but in *T. brucei* and *Leishmania* both BDFs coprecipitate as members of CRKT [19, 20]. Here, we confirmed the extra-nuclear localization of BDF1 and BDF3, but they also appear in our CRKT interaction network, which we identified as a mix of nuclear and non-nuclear components. This apparent contradiction suggests that BDFs may shuttle between compartments, potentially linking cytoplasmic processes with nuclear regulation. In the case of BDF3, its mobilization to the flagellum during metacyclogenesis links it to a fundamental event in the life cycle of *T. cruzi* [15]. On the other hand, BDF1 has been linked with glycosomes [14]. Notably, the identification of the αN domain of BDF1 as a peroxisomal localization signal type 2 (PTS-2), and the fact that the TINB1 domain of BDF4 binds to this domain, may suggest a cross-interaction between energy metabolism and epigenetic regulation. We propose that BDF4 recruits BDF1 to the nucleus by blocking the PTS-2 in specific metabolic situations, ultimately regulating nuclear events. While much research is needed to confirm this hypothesis, there are indications that such mechanisms could also be present in other trypanosomatids. In *T. brucei*, the subcellular localization of BDF1 observed in the TrypTag database varies depending on the position of the tag used; the N-terminal tag, which would block a potential peptide transit, is nuclear, but the C-terminal tag shows a localization consistent with glycosomes [29]. On the other hand, Staneva et al. observed a nuclear-cytoplasmic localization for BDF4 [17].

We also found that the ET domain of BDF3 may recognize linear motifs in BDF4, BDF5 and EPL2ap, with interaction modes similar to those of other ET domains [62]. These predicted dynamic interactions may accelerate the recruitment of these proteins to the CRKT complex or help position them in appropriate orientations. Further exploration of the interactions mediated by the ET domain of BDF1 outside the context of CRKT is still required.

Our BDF6 proximity proteome recovered several identifiable NuA4 components with high enrichment, consistent with co-IP assays performed in *T. brucei* [18, 19]. Among the nine proteins recovered in both *T. cruzi* (this study) and *T. brucei*, two corresponded to unknown proteins that we successfully assigned to conserved protein families. One was identified as a member of the ING protein family (INGL). The other was structurally similar to the N-terminal domain of DMAP1 (DNTL). Members of both families are NuA4 components in other organisms [41, 42]. The presence of a NuA4 complex in trypanosomatids has important evolutionary implications. Although many of the identified components share limited sequence identity with their homologs in Opisthokonta, the overall structure of the complex exhibits a striking similarity to yeast piccolo-NuA4. An analogous situation was previously observed for the TINTIN subcomplex [19]. Given that Discoba is considered one of the earliest diverging phylogenetic groups, our results suggest that the emergence of a NuA4-nucleosome mode of interaction is an early event in the evolution of eukaryotes. Considering that the catalytic activity associated with NuA4 is the only essential HAT activity in yeast reinforces this hypothesis [65].

In Opisthokonta, NuA4 is dynamic, shares subunits with other complexes, and is composed of multiple modules with DNA repair and transcription functions [66]. Piccolo-NuA4 can function as an independent nucleosome-directed HAT module responsible for widespread basal acetylation, whereas the full 13-subunit NuA4, which includes the TRA (<u>Tr</u>anscription <u>A</u>ctivator-binding) module, enables targeted acetylation through transcription factor-mediated recruitment to promoters [67]. The recruitment of TRA is mediated through the EPcC domain of EPL1, a piccolo-NuA4 subunit [65]. We found that trypanosomatid EPL1 lacks this domain, while the TRA module is completely absent. A version of NuA4 closer to piccolo than to the complete complex is consistent with the fact that trypanosomatid protein-coding genes are transcribed polycistronically and lack gene-specific RNA Pol II promoters. On the other hand, yeast TINTIN (EAF3, EAF5 and EAF7) constitutes a functionally distinct module that operates both within the full complex and independently to couple NuA4 activity to transcription elongation, helping recruit the entire NuA4 complex to chromatin. This targeting depends on EAF3’s (MRGx homolog) chromodomain (CD), which recognizes H3K36 methylation, thereby increasing NuA4 occupancy across transcribed gene bodies and promoting H4 acetylation during elongation [22]. Since this domain is absent in the trypanosomatid version of TINTIN, and NuA4 resembles a piccolo-NuA4 rather than the full-size complex, we can expect mechanistic differences from its homologous complexes in Opisthokonta. For example, the BD of BDF6 may act as a functional substitute for the CD, recognizing acetylation signals instead of methylations, as the BD of EAF5 (the BDF6 homolog) is non-functional in yeast [21]. Furthermore, considering that the Tudor domain of *T. brucei* HAT1 is unable to recognize methylated lysines (Kme) on histone tails and instead exhibits nonspecific DNA binding capacity [48], something similar may occur in *T. cruzi*. Both observations suggest that the recognition of Kme is not as important for the function of these complexes as it is in other organisms.

Lastly, both the predicted interaction map and the proximity network indicate cross-interactions between CRKT and NuA4. This is consistent with previous observations in *T. brucei*, where multiple BDF proteins co-occupy transcription start regions (TSRs) [19, 68]. More broadly, interactions between epigenetic complexes are common in eukaryotes, as many of these assemblies share subunits and form modular, partially overlapping networks. For example, components such as RuvBL1/2 are found in multiple chromatin remodeling and acetyltransferase complexes, and MRG family proteins participate in distinct regulatory assemblies, highlighting the extensive reuse of core factors across complexes [57, 66]. Beyond the BD-mediated recognition of acetylated nucleosomes, we propose that their positioning at TSRs is also regulated by dynamic interactions between different modules. For example, chromatin-bound BDF5 would engage EPL1, thereby recruiting it to TSRs. Once HAT1 binds EPL1, BDF5 would release EPL1, seeding the assembly of the catalytic module of NuA4 and the recruitment of BDF6. NuA4 modules would also be able to form independently and recruit BDF5 when EPL1 is not yet bound to HAT1. This mechanism could propagate acetylation marks between adjacent nucleosomes through a “walking EPL1” and provide a molecular route for the maintenance and coordination of heritable acetylation patterns at TSRs. Moreover, a multibromodomain complex with symmetric architecture like the CRKT core would likely amplify this mechanism.

In summary, our results provide valuable structural and functional information about two trypanosomatid bromodomain-containing complexes that integrate nuclear and cytoplasmic components (**Fig. *7***). Although our structural modeling approaches do not replace experimental validation, they provide a reliable framework to generate testable hypotheses. Moreover, they also enable the prediction of dynamic states arising from the promiscuity of certain components, which are often difficult to capture using biophysical methods because they may represent low-population conformations within the broader conformational landscape. Through a rapid and low-cost computational strategy, our method adds a structural dimension to flat interactomics data that allows for the prediction of key functional interactions that may be suitable for intervention. For example, the BDF5–BDF5BP interaction, previously shown to be susceptible to disruption [21], in now revealed as essential for CRKT integrity, since its loss disconnects the PARPL–BDF8–BDF5 arm from the core module. Similarly, the engagement of the CTD of BDF6 with EPL1 is the physical link between TINTIN and the catalytic module of NuA4, and is therefore critical for maintaining functional coupling between these modules. Both interactions can be tested as drug targets by designing screening systems for compounds that block these surfaces. By revealing these interfaces, we provide a structural basis for targeting critical protein–protein interfaces, enabling the development of trypanocidal molecules with greater specificity than the established BD-pocket inhibition paradigm.

**Fig. 7:**
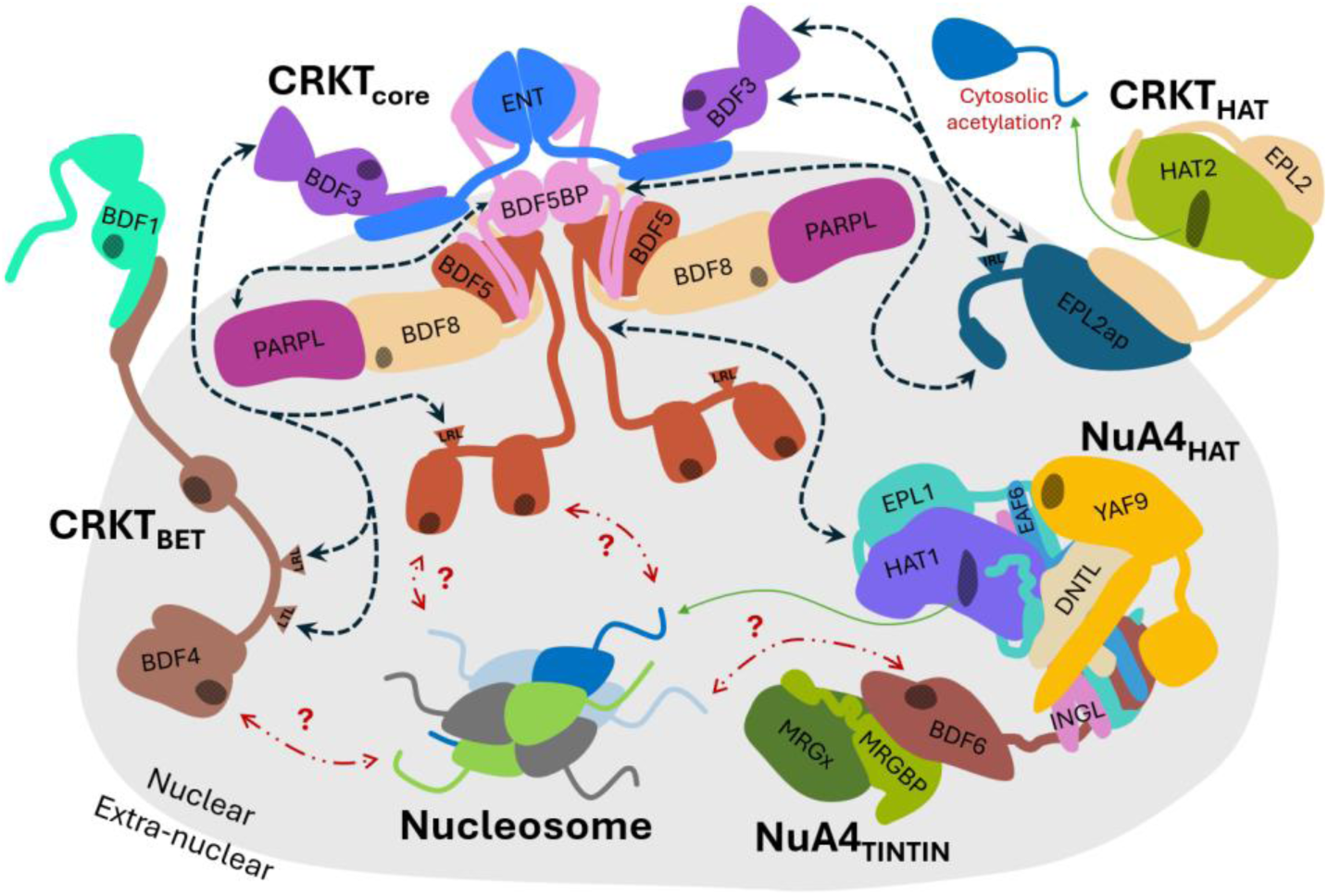
Model summary of NuA4 and CRKT modules from *T. cruzi*. The nuclear confidence level of each protein is represented by being inside or outside of the gray background. Predicted dynamic interactions are highlighted with black dashed arrows. Catalytic activities are represented with green arrows and red dashed arrows indicate unknown specificity for acetylated residues.

## Materials and Methods

### pTcTurboID design and coding sequences cloning

The pTcTurboID vector was designed based on the pTcINDEX-GW vector, previously optimized in our laboratory for tetracycline-regulated expression in *T. cruzi* [69]. To this end, the 3xHA-TurboID cassette from the 3xHA-TurboID-NLS_pCDNA3 vector (<u>Addgene</u> <u>#107171</u>) was PCR-amplified using the 3xHA-TurboID-Fw and 3xHA-TurboID-Rv oligos (**Supplementary Table *3***). The cassette was then inserted into the ClaI restriction site of the pTcINDEX-GW vector (<u>Addgene #234713</u>), downstream and in phase with the Gateway cassette, resulting in the pTcTurboID-GW vector (<u>Addgene #234714</u>). Proper orientation was verified by PCR and the expected sequence was confirmed by Sanger sequencing.

The sequences of BDF2, BDF3, BDF4, BDF5, BDF6, BDF8, and GFP were cloned into the Gateway pENTR3c entry vector using the corresponding oligos and restriction enzymes from **Supplementary Table *3***. As templates, *T. cruzi* Dm28c genomic DNA and an entry plasmid containing the GFP sequence mgfp5 were used, respectively. An additional cassette containing three nuclear localization signals (3xNLS) from the SV40 virus was extracted from the 3xHA-TurboID-NLS_pCDNA3 vector using the restriction enzymes EcoRI and XhoI and inserted into the pENTR3c vector using the same enzymes. From the resulting vector, a 233 bp fragment containing the 3xNLS cassette was extracted using the enzymes HincII and PstI and then introduced in frame with the CDS of GFP into the pENTR3c_GFP vector cut with EcoRV and PstI (2870 bp fragment). The expected sequences of all entry vectors were confirmed by Sanger sequencing.

Finally, the CDSs of GFP, GFP-3xNLS, BDF2, BDF3, BDF4, BDF5, BDF6, and BDF8 were transferred from the pENTR3c vectors to pTcTurboID via LR recombination [70]. The resulting vectors from the LR cloning of GFP (<u>Addgene #234715</u>) and GFP-3xNLS (<u>Addgene #234716</u>) are available from Addgene.

### Culture, transfection and direct cloning of epimastigotes

Epimastigotes of the Dm28c pLew13 strain were cultured in LIT medium supplemented with 200 μg/mL of G418 and maintained in exponential phase for two weeks (10–30 × 10⁶ epimastigotes/mL). The culture was then expanded to reach the required number of parasites. For each transfection, 60–80 μg of circular pTcTurboID plasmids or 15–30 μg of SpeI-linearized plasmid were precipitated with isopropanol and resuspended in 25 μL of TbBSF transfection buffer [71]. 40 x 10⁶ epimastigotes were washed with 1.5 mL of PBS, resuspended in 375 μL of TbBSF buffer, and gently mixed with the 25 μL of plasmid by pipetting. The mixture was transferred to an electroporation cubette and incubated on ice for 10 minutes. A negative control using only transfection buffer was added at each transfection round. Nucleofection was performed using a single pulse of the X-014 program on the Amaxa Nucleofector 2b electroporator (Lonza Cologne AG, Germany). The parasites were immediately resuspended in 12 mL of antibiotic-free supplemented LIT medium and incubated in T-25 bottles at 28 °C.

After 24 hours, G418 (200 μg/mL) and hygromycin (200 μg/mL) were added. The culture was homogenized, and selection and direct cloning were performed in 24-well plates, distributing 500 μL of culture per well. The plates were incubated at 28 °C in airtight containers with a humid chamber, and the cultures were observed regularly. After 4 to 8 weeks, 1 to 7 wells with actively growing epimastigotes were obtained per transfection [26]. Up to 3 positive cultures per transfection were randomly selected and expanded in LIT medium supplemented with G418 (200 μg/mL) and hygromycin (200 μg/mL) to confirm plasmid integration and expression of the selected genes by PCR and western blot, respectively.

### Western blots

Between 30 and 50 μg of protein extracts were fractionated by 12% SDS-PAGEs and transferred to nitrocellulose membranes. The transferred proteins were visualized using Ponceau S staining. The membranes were blocked with 10% skim milk in PBS for 2 hours, rinsed three times with PBS containing 0.5% Tween20 (PBS-T), and then incubated with specific antibodies/conjugates. To visualize the bands corresponding to fusions, the membranes were incubated with 0.1 μg/mL rat anti-HA (Roche) resuspended in PBS-T for 3 hours and then rinsed three times with PBS-T. Bound antibodies were detected using rabbit anti-rat IgG secondary antibodies labeled with horseradish peroxidase (GE Healthcare). To visualize biotinylated proteins, membranes were incubated with 0.3 μg/mL of peroxidase-conjugated streptavidin (Jackson ImmunoResearch) resuspended in 0.1% PBS-T and 3% BSA for 1 hour at room temperature. To reveal peroxidase activity, membranes were washed three times with PBS-T and visualized using the ECL Prime kit (GE Healthcare) according to the manufacturer protocol. Immunoreactive bands were visualized on an Amersham Imager 600 digital imaging system. In some assays, the amount of protein extract loaded was verified by fractionating an SDS-PAGE identical to the one transferred, which was then stained with Coomassie Blue.

### Indirect immunofluorescence

For each condition, 10 × 10⁶ epimastigotes were taken and washed three times with PBS, centrifuging at 1500 g for 5 minutes, and discarding the supernatant between each wash. The parasites were resuspended in 100 μL of 4% p-formaldehyde in PBS and 20 μL were loaded per well, incubating for 20 minutes on immunofluorescence slides previously treated with poly-lysine. After three washes with PBS, epimastigotes were permeabilized with 30 μL of 0.2% Triton X-100 in PBS for 10 minutes and washed again three times with PBS. The parasites were incubated for 3 hours in a humid chamber with 30 μL of primary anti-HA antibody (BSA 1%, Roche rat anti-HA 1 μg/ml, PBS) and washed three times with 0.01% PBS Tween-20. Subsequently, they were incubated with 30 μL of FITC-conjugated goat anti-rat secondary antibody (Life Technologies) at a concentration of 1 μg/mL and 2 μg/mL of 4′,6-diamidino-2-phenylindole (DAPI) in PBS for 1 hour. Three final washes were performed with 0.01% PBS Tween-20, and the slide was mounted using Vectashield®. Images were acquired using a Nikon Eclipse Ni-U fluorescence microscope.

### Purification with streptavidin magnetic beads

#### Setup

For each condition, culture medium containing 150 × 10⁶ epimastigotes induced for 24 h with the chosen tetracycline concentration was separated and cooled on ice for 15 minutes. They were then harvested, washed with cold PBS and resuspended in 750 μL of RIPA buffer (50 mM Tris, 150 mM NaCl, 0.1% SDS, 0.5% sodium deoxycholate, 1% Triton X-100) supplemented with protease inhibitors (AEBSF 1 mM, Aprotinin 2 µg/ml, Bestatin 1 µM, E-64 10 µM, Leupeptin 10 µM, Pepstatin A 1 µM and PMSF 1 mM final). After 30 minutes on ice, the samples were sonicated in two 10-minute cycles (30 sec ON, 30 sec OFF, Mid power) using a Bioruptor® sonicator. Each sample was divided into two portions: 500 µL (100 x 10⁶ cells/mL) and 250 µL (50 x 10⁶ cells/mL). The 250 µL portion was stored at −20°C for later analysis (INPUT). The remaining portion was centrifuged at 20,000 g for 10 minutes at 4°C, and the supernatant was recovered. 25 µL of magnetic beads (Pierce™) were prepared and washed twice with 1 mL of RIPA buffer. All washes were performed on a magnetic rack unless otherwise noted. The supernatant from the 500 µL was mixed with the washed beads and incubated for 16 hours at 4°C on rotary tube mixer. The supernatant was recovered for further analysis (FLOW THROUGH). Subsequently, sequential washes were performed with 1 mL of RIPA (twice), 1 mL of 1 M KCl, 1 mL of 0.1 M CaCO₃, 1 mL of 2 M urea in 10 mM Tris pH 8, and 1 mL of RIPA (twice), transferring the beads to new tubes during the final wash. Finally, the beads were resuspended in 30 µL of cracking buffer (3X loading buffer, 2 mM biotin, and 20 mM DTT) and incubated for 10 minutes at 95°C. The resulting supernatant was stored at −20°C (ELUATES).

#### Mass spectrometry

For mass spectrometry analysis, 400 × 10⁶ epimastigotes induced for 24 h in fresh LIT medium with the optimum tetracycline concentration were harvested per replicate. The parasites were processed following the protocol described for the setup, with the following modifications: the entire volume of parasites was used in a single tube with 500 µL of RIPA (without separating fractions for SDS-PAGE analysis), and 100 µL of beads were used instead of 25 µL per replicate. After the final wash with RIPA buffer (described for the setup), the beads were further washed sequentially with 1 mL of 50 mM Tris-HCl pH 7.5 and twice with 1 mL of 2M urea in 50 mM Tris-HCl pH 7.5. During the final wash, the beads were transferred to new tubes. The proteins were digested directly on the beads (on-bead digestion) for 16 h at 37°C under horizontal orbital shaking (1200 rpm) with 200 μL of digestion buffer (2M Urea, 50 mM Tris-HCl pH 7.5, 1 mM CaCl2, 2 mM DTT) containing 0.5 μg of trypsin. The supernatant was collected in a new tube, and the beads were rinsed with an additional 100 μL of 2M Urea, 50 mM Tris-HCl, pH 7.5 buffer, which was then combined with the first supernatant. All conditions were analyzed in quadruplicate.

### Silver staining

Eluates obtained during setup were heated to 95°C for 10 minutes and centrifuged at 12,000 g for 3 minutes. The proteins in the resulting supernatant were then separated by 12% SDS-PAGE, using 10 µL per well. The gel was then immersed in 50 mL of fixation solution (50% methanol; 12% acetic acid; 0.05% formaldehyde), heated for 30 seconds in a microwave oven at full power, and incubated for 5 minutes with shaking at room temperature. Subsequently, sequential incubations were performed with 50 mL of 30% ethanol and then with 50 mL of 0.02% sodium thiosulfate, each for 5 minutes with shaking at room temperature and an initial heating of 30 seconds. Two washes were performed with H_2_O miliQ (30 seconds of heating, 2 minutes of shaking), and the gel was then incubated with 30 mL of silver solution (0.2% AgNO3; 0.075% formaldehyde) for 5 minutes with shaking at room temperature, after a 30-second heating. After a wash with H2O miliQ (20–60 seconds) at room temperature, the gel was developed by incubation with developing solution (6% Na_2_CO_3_; 0.05% formaldehyde; 0.0004% sodium thiosulfate) for 5–10 minutes at room temperature. The reaction was stopped by adding stop solution (50% methanol, 12% acetic acid). All solutions were prepared fresh using H_2_O miliQ.

### Mass spectrometry coupled with liquid chromatography

Peptides obtained by on-bead digestion with trypsin were treated with iodoacetamide (10 mM final concentration) for 20 minutes and concentrated by evaporation in a SpeedVac to a volume of 100 μL. The samples were desalted using C18 resins (ZipTip C18, Millipore), dried again by evaporation, and resuspended in solvent A (0.1% formic acid in water, Milli-Q). Liquid chromatography-mass spectrometry (LC-MS) analysis was performed on an Ultimate3000 nanoHPLC system (Thermo Scientific) equipped with a 25 cm² nano C18 column (ES902, Thermo Fisher Scientific). 3 μL of each sample were injected, and separation was performed using 0.1% formic acid in water (solvent A) and 0.1% formic acid in acetonitrile (solvent B). The gradient consisted of 4–30% solvent B for 64 minutes, followed by 30–80% solvent B for 7 minutes and 80% solvent B for 1 minute. Mass spectrometry analysis was performed on a Q-Exactive HF (Thermo Scientific) operating in positive mode, with a liquid junction voltage of 1.9 kV and a capillary temperature of 300°C. Full spectra were acquired in the m/z range of 375–2000 with a resolution of 120,000 (at m/z 200), automatic gain control (AGC) of 1 × 10⁶, and a maximum injection time of 100 ms. The 10 most intense precursor ions were selected for MS/MS fragmentation using a normalized collision energy of 27 eV. MS/MS spectra were acquired with first dynamic mass, AGC of 5 × 10^5^, resolution of 30,000 (at m/z 200), isolation window of 1.4 m/z and maximum injection time of 55 ms. Ions with no assigned charge, single charge or greater than 6 were excluded, implementing a dynamic exclusion of 25 seconds.

### Structure prediction and analysis

Protein structure prediction and analysis using AlphaFold3 (AF3) and MultimerMapper (MM), respectively, was conducted as described previously [26], with small modifications. Briefly, a FASTA file containing the protein sequences of each system (**Table 2** and **Table 1**) was created. Each file was loaded into MM v1.0.0 [24] to initialize the systems, i.e., get all dimeric suggestions (2-mers), including homodimers. The 2-mers were then predicted using AF3 server and resulting structures were converted to a compatible format using the af3_compatibilty.py utility from MM. Converted data was fed into MM to detect interactors and suggest 3-mers based on interacting pairs. 3-mers were predicted with AF3, converted and fed back with the 2-mers to MM to generate new suggestions. The process was repeated iteratively to explore available branches of the stoichiometric space and find convergent stoichiometries, i.e., protein combinations in which all possible children were unstable. MM cutoffs used to determine stable complexes, interactions and contacts were 8.3 Å (minimum interaction PAE) and 4 (N° of models).

### Combinatorial Assembly

CombFold was used to perform the combinatorial assembly of the CRKT core module [94]. Protein subunits were defined based on the domains detected with the domain detection algorithm for BDF3, BDF5, ENT, BDF5BP, BDF8 and PARPL, excluding disordered segments (**Fig. *5*A**). Subsequently, a subunits.json file was generated where each subunit was defined by its name, its amino acid sequence, its associated chains, and the start residues within the original sequence. The number of subunits used for each protein domain or protein was set as two.

For the assembly, a folder was prepared containing all the structural predictions for the subunits to be assembled in PDB format (pdbs directory) and the subunits.json file. It was executed using the following command:

CombFold/scripts/run_on_pdbs.py subunits.json pdbs out

The resulting assembled structures within the out directory were subsequently analyzed and visualized to evaluate the assembly quality and the biological validity of the models. Substructures that were not in contact with the main core were reoriented for better visualization: the bromodomains of BDF5 and the FHA-ENT and BDF3-TINB3 substructures.

## Supporting information

supp_file_1-Experimental_round_1_ProteomeDiscoverer

supp_file_2-Experimental_round_2_ProteomeDiscoverer

supp_file_3-annotated_nuclear_proximity_proteome

supp_file_4-proteins_classificated_as_proximal_or_interactors

supp_file_5-symbols_used_in_network

## Acknowledgments

This work was funded by the National Research Council (CONICET: PIP 2021-0848); the National Agency for Science (ANPCyT: 2020-01704); Universidad Nacional de Rosario (UNR: 0001-00285485); Agencia Santafesina de Ciencia, Tecnología e Innovación, Santa Fe, Argentina (PEICID2023-085) and AWS Cloud Credit for Research Program (PS_R_FY2022_Q2_CONICET_Serra_10900). We want to thanks Alejo Cantoia and Ines Espindola for their technical assistance and Tess C. Branon for sharing valuable methodological information about TurboID-proximity labeling experimental setup.

