## Supplementary figures and tables for "Structural Insights into Bromodomain-Containing Complexes from *Trypanosoma cruzi* Revealed by Proximity Labeling and Stoichiometric Space Exploration"

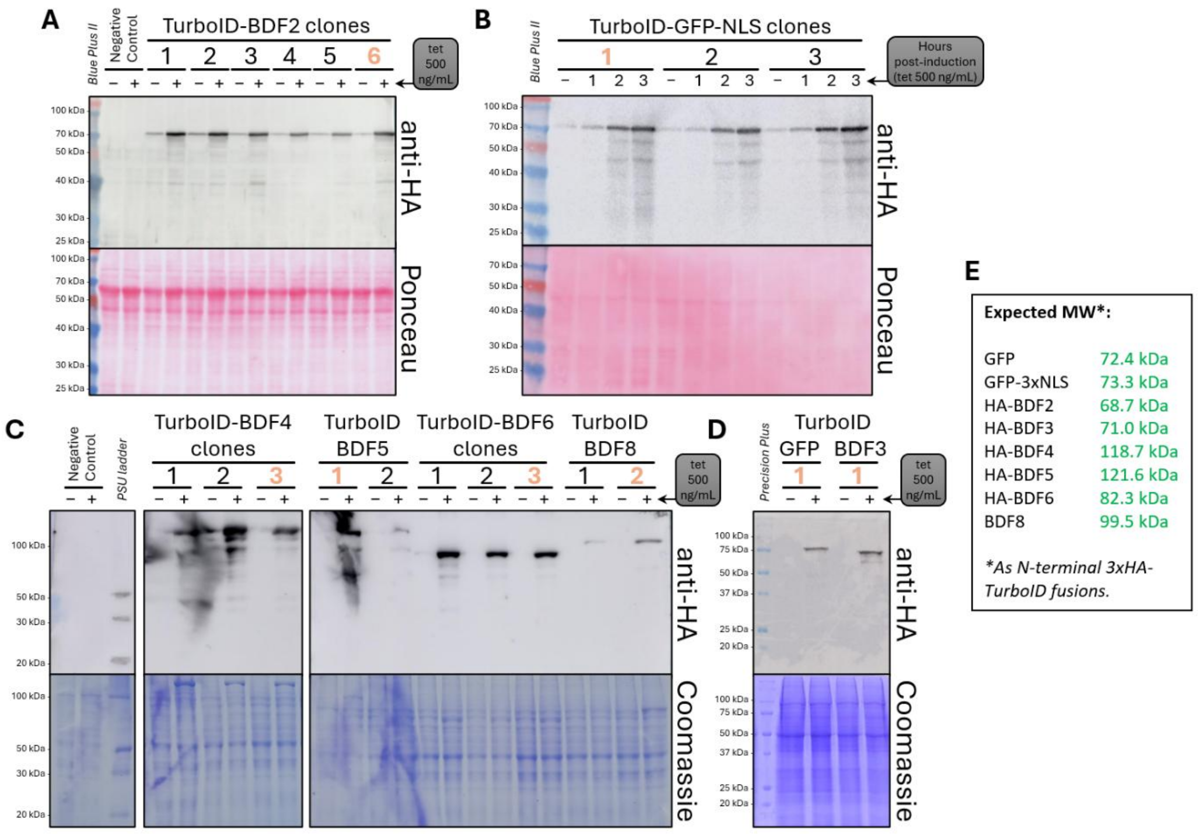


**Supplementary Fig. 1: Induction test of clones expressing TurboID fusions using maximum expression tetracycline concentration**. Clones were induced by 24 hours using high tetracycline (tet) levels (500 ng/mL), which maximize the expression of the transgenes (+), and were compared to the uninduced condition (–). For TurboID-GFP-NLS, the same concentration was used, but inducing for different times (1, 2 and 3 hours). Selected clones for further studies are highlighted in orange. Transgene expression for the 6 clones obtained of TurboID-BDF2 (**A**), 3 clones of TurboID-GFP-NLS (**B**), 3 clones of TurboID-BDF4, 2 clones of TurboID-BDF5, 3 clones of TurboID-BDF6, 2 clones of TurboID-BDF8 (**C**), 1 clone for TurboID-GFP and 1 clone for TurboID-BDF3 (**D**). (**E**) Expected molecular weights of N-terminal tagged 3xHA-TurboID fusions.

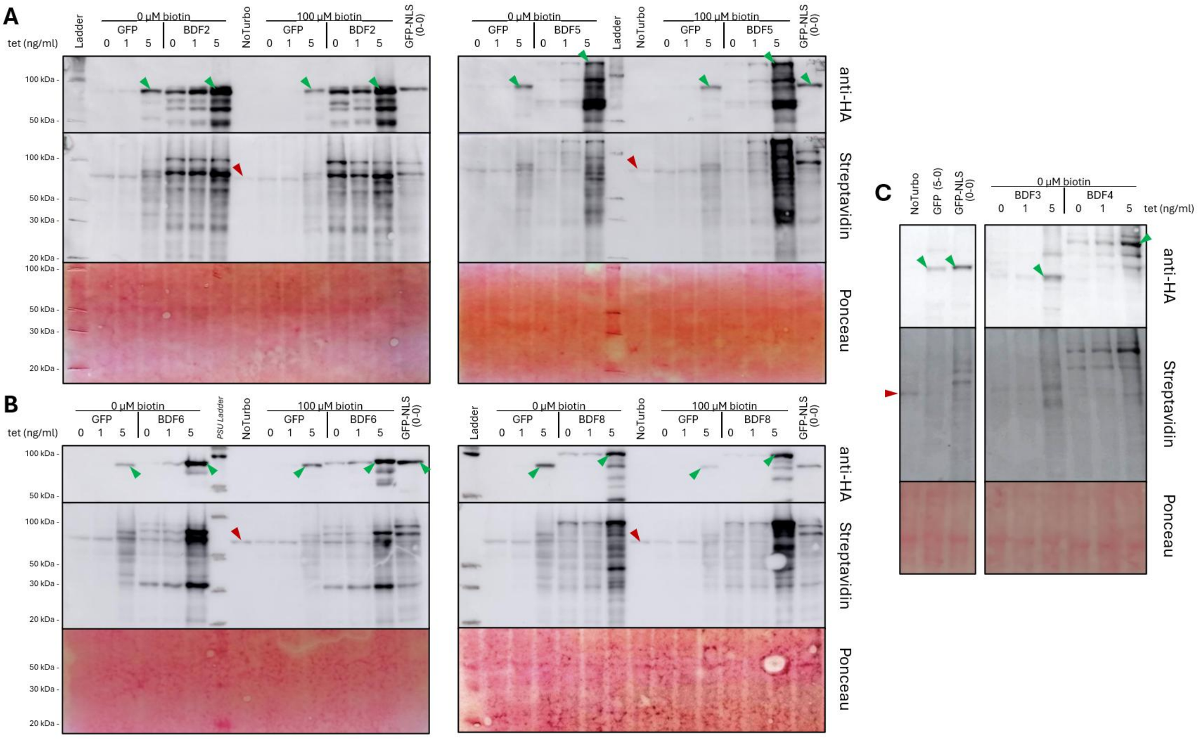


**Supplementary Fig. 2: Expression levels normalization using the naturally biotinylated 3MCC band as reference.** Western blots showing the expression of 3xHA-TurboID fusions using anti-HA and biotinylation patterns using streptavidin-HRP in response to tetracycline and 100 µM biotin supplementation for clones expressing GFP, GFP-NLS, BDF2, and BDF5 (**A**); and GFP, GFP-NLS, BDF6 and BDF8 (**B**). (**C**) Western blots for GFP, GFP-NLS, BDF3 and BDF4. Green arrows point to the bands corresponding to the fusion proteins, while red arrows point to the naturally biotinylated 3MCC band in the control without TurboID (No Turbo). Ponceau stain was used to verify even load. Epimastigotes clones were incubated for 24 h at 28ºC with different concentrations of tetracycline and biotin in fresh supplemented LIT medium.

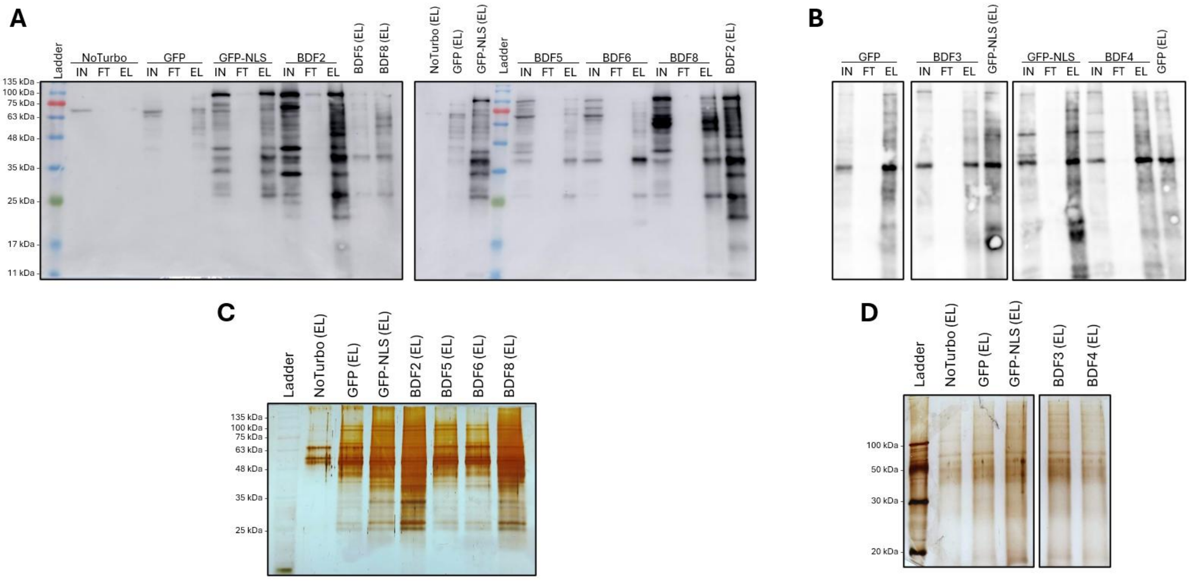


**Supplementary Fig. 3: Western blot analysis with streptavidin-HRP and silver staining of the purification process**. (**A** and **B**) Different fractions of the purification process using magnetic beads for each linage, analyzed with streptavidin-HRP to detect biotinylated proteins. (**A**) First purification test. All eluates are present on both membranes for comparison. (**B**) Second purification test. Silver staining of the eluates from the first (**C**) and second (**D**) purification tests. **NoTurbo**: Negative control linage without TurboID. **IN**: Input; **FT**: Flow-Through; **EL**: Eluate.

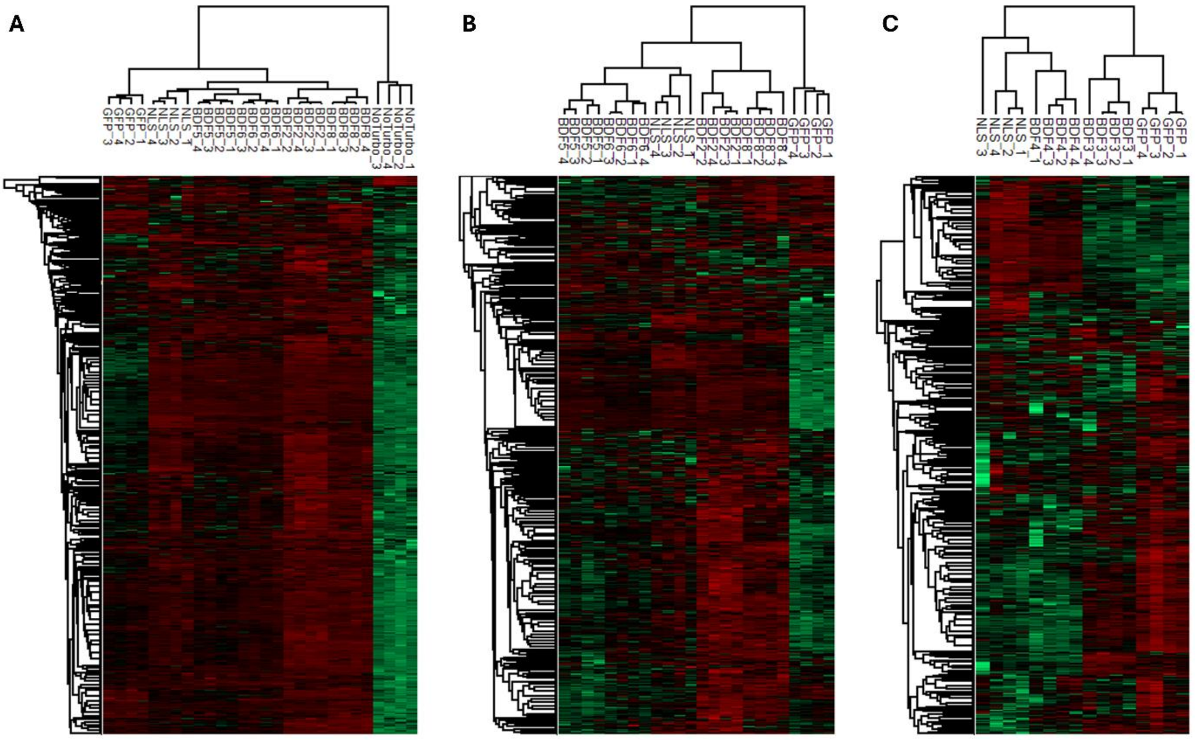


**Supplementary Fig. 4: Clustering analysis of samples and proteins using heat maps**. Heat maps using hierarchical clustering at the protein (rows) and sample (columns) levels. Normalized intensities using the Z-score metric were used. The most abundant proteins are shown in red and the least abundant in green. Replicates of each condition are indicated as *condition_i*. (**A**) Results using all samples from the first experimental round. (**B**) Results from the first experimental round removing the control sample (NoTurbo) and normalizing again using the Z-score method. (**C**) Results for all samples from the second experimental round.

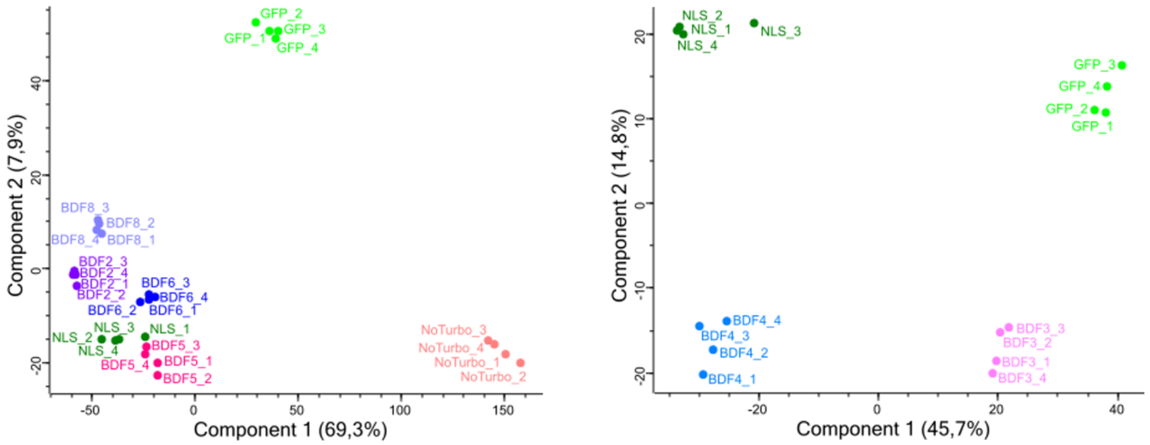


**Supplementary Fig. 5: Principal component analysis (PCA)**. PCA analysis for experimental rounds 1 (**left**) and 2 (**right**). The nomenclature used is that of **Supplementary Table 1**, indicating the replicate number with an underscore and a number.

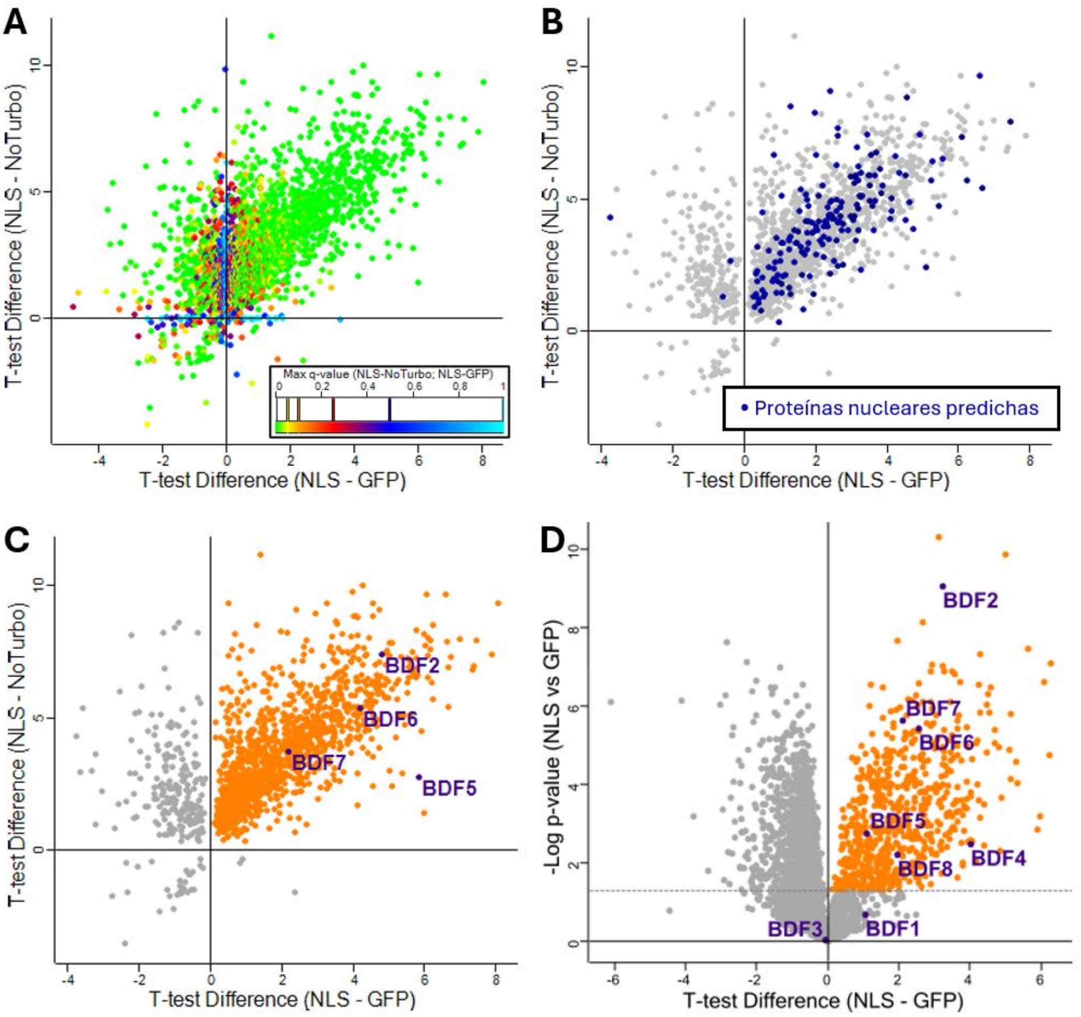


**Supplementary Fig. 6: Nuclear proximity proteomes from two different experiments**. Each detected protein is represented by a point in the plots. (**A**) Scatter plot comparing the difference in the T-test between the signal for each protein in the nuclear and cytoplasmic spatial controls (NLS – GFP) versus the signal between the nuclear spatial control and the control without TurboID (NLS – NoTurbo). The color scale represents the maximum q-value (FDR) of both tests for each protein. (**B**) Same plot as in panel A, but filtering at a significance level of FDR=0.01. Proteins colored blue are all those containing the words “nucleus,” “nuclear,” “nucleolus,” or “chromosome” in any of their annotations. (**C**) Same graph as in panel A, but filtering at a significance level of FDR=0.01. The proteins colored orange are those that were significantly enriched in the nuclear spatial control compared to both the cytoplasmic spatial control and the control without TurboID. BDFs detected in the samples that passed the filter are indicated in purple. (**D**) Volcano plot for the comparison of the nuclear and cytoplasmic spatial controls of the second experimental round. The orange proteins are those that were significantly enriched in the nuclear spatial control compared to the cytoplasmic spatial control (FDR=0.05). BDFs are indicated in purple.

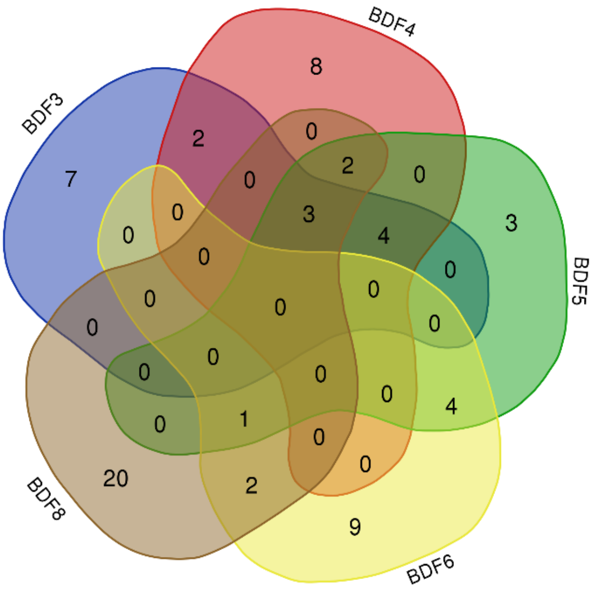


**Supplementary Fig. 7: Interactors overlap analysis.** Venn diagram showing the distribution of interacting proteins among the different PL experiments. Only proteins classified as "interactors" according to the established criteria were considered. The diagram reveals a core of proteins shared among BDF3, BDF4, BDF5, and BDF8, while BDF6 shares only some proteins with BDF5 and BDF8.

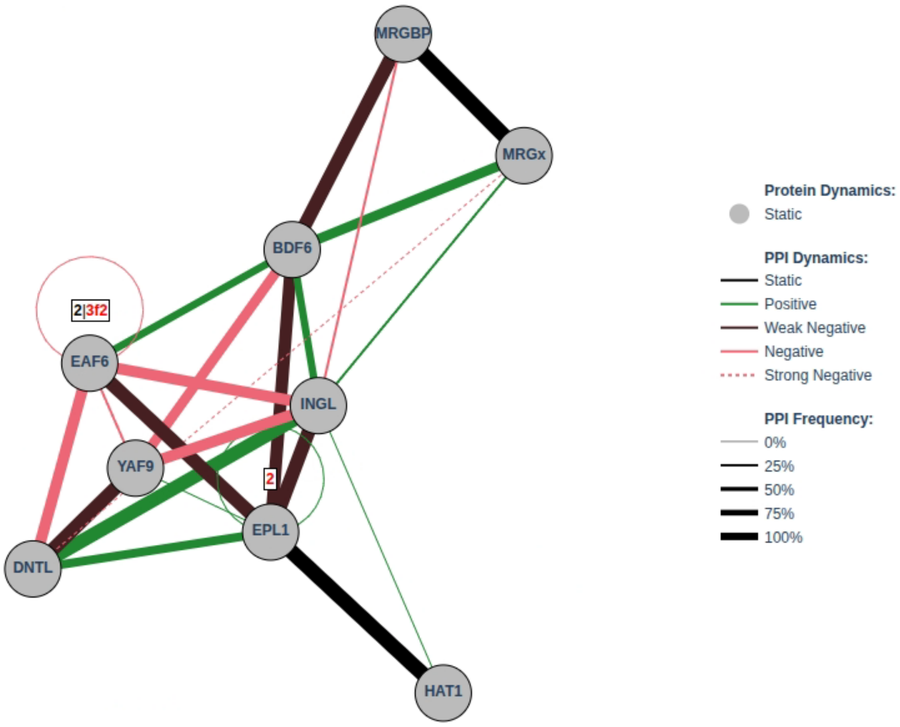


**Supplementary Fig. 8: Full predicted PPI graph of NuA4 components obtained with MultimerMapper**. Extra interactions compared to **Error! Reference source not found.**C are, for example, those PPIs that are induced when another interaction causes two proteins to be closer in space and get in contact (positive dynamic interactions). For example, BDF6 is unable to bind MRGx in 2-mer predictions, but they interact when MRGBP is present (3-mer). This results in a positive dynamic interaction between BDF6 and MRGx, activated by MRGBP.

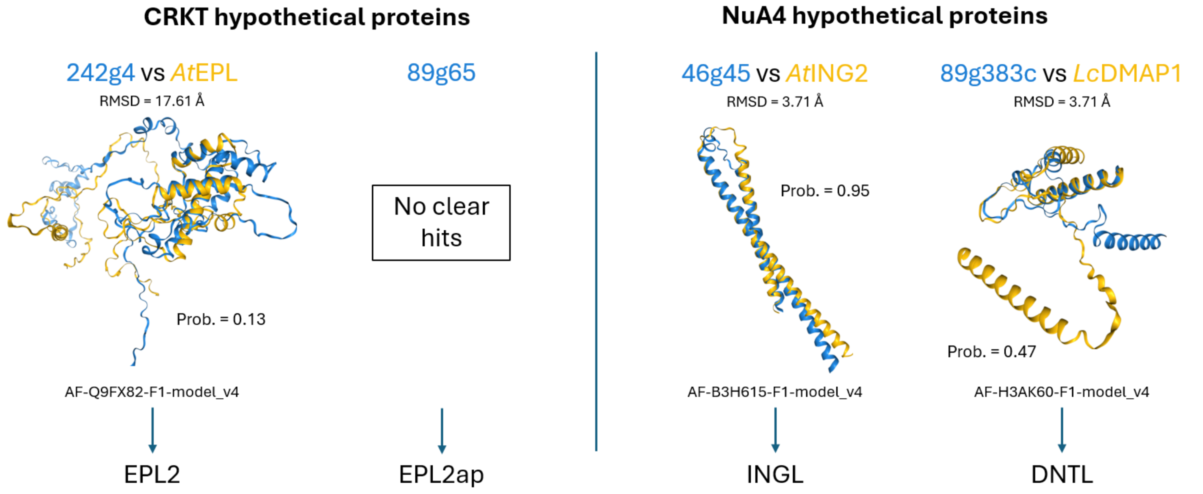


**Supplementary Fig. 9: Foldseek structural alignments of the hypothetical proteins from CRKT and NuA4 of *T. cruzi*.** The backbone of the query proteins is shown in blue, and that of the most significant hit is shown in yellow. The symbol chosen to represent each protein is shown below. **EPL**: Enhancer of Polycomb-Like. **EPL2ap**: EPL2-associated protein. **INGL**: ING-Like. **DNTL**: DMAP1 N-Terminal Like.

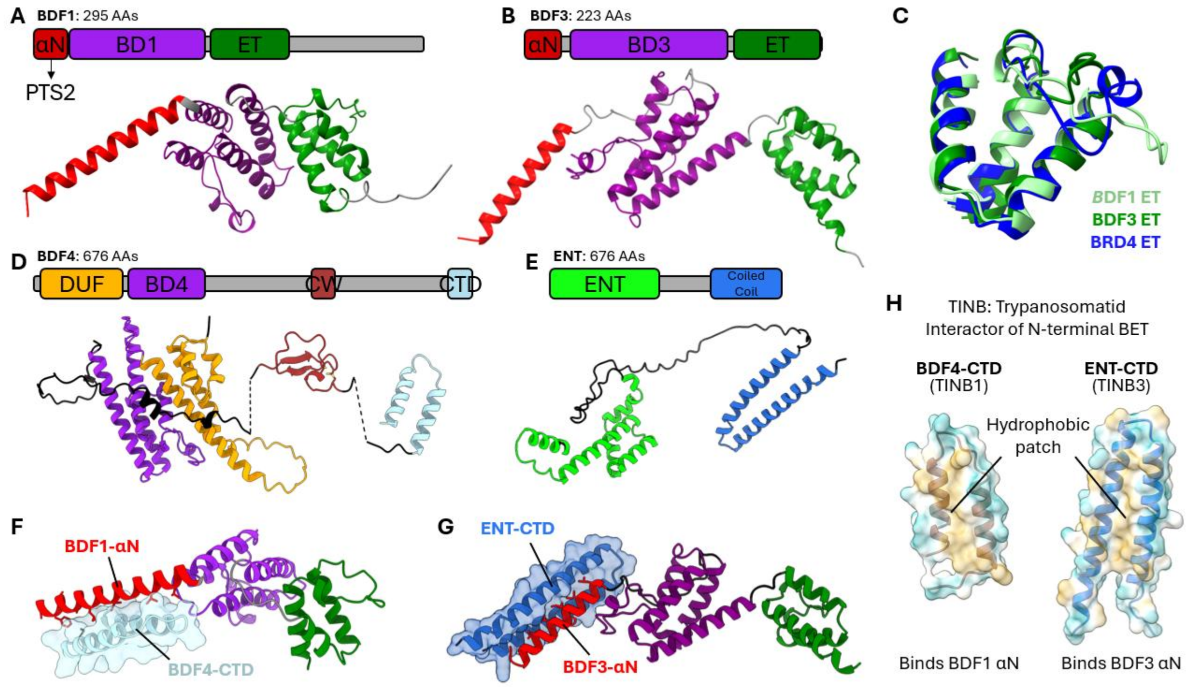


**Supplementary Fig. 10: Domains of the *T. cruzi* proteins BDF1, BDF3, BDF4, and ENT and their interaction modes predicted by MultimerMapper**. (**A**) BDF1 and (**B**) BDF3 domains. (**C**) ET domain of BDF4 structurally aligned with the C-terminal domains of BDF1 and BDF3. Domain architecture of BDF4 (**D**) and ENT (**E**). Representative structures of the interaction modes between BDF1-BDF4 (**F**) and BDF3-ENT (**G**) pairs. (**H**) Solvent-exposed surface of the CTD domains of BDF4 and ENT, colored by hydrophobic potential. Proposed names for these domains are indicated in parentheses. **αN**: N-terminal α-helix domain; **BD**: Bromodomain; **ET**: Extra-Terminal domain. **DUF**: Domain of Unknown Function. **ENT**: EMSY N-Terminal.

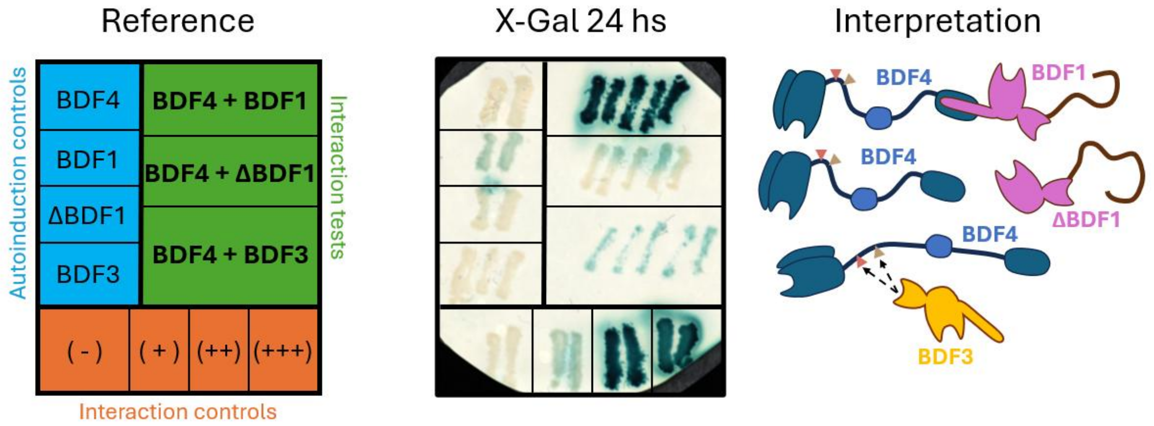


**Supplementary Fig. 11: Interaction tests between BDF1-BDF4 and BDF3-BDF4 pairs using Y2H.** Mav203 yeast strain was transformed with pGADT7 and pGBKT7 vectors to express different versions of BDF1, BDF3, and BDF4. The reference (**left**) indicates the genes that were expressed in the yeasts and the interaction strength of the control strains. Colonies were randomly selected and streaked onto a plate containing YPAD medium with a sterile nitrocellulose membrane on top. The membrane was removed and incubated for 24 h in X-gal medium to detect β-galactosidase expression, the levels of which are directly proportional to the interaction strength (**center**). The interpretation of the results is shown on the **right**.

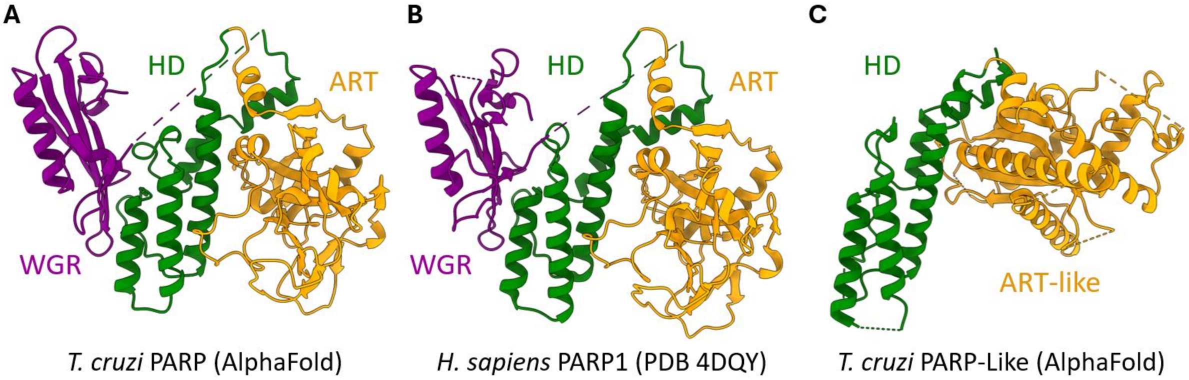


**Supplementary Fig. 12: Structural comparison between the PARP and PARPL proteins of *T. cruzi* with human PARP1**. (**A**) AF2 model of PARP (*T. cruzi*). (**B**) X-ray crystal model (PDB 4DQY) of PARP1 (*H. sapiens*). (**C**) AF3 model of PARPL (PARP-like) from *T. cruzi*.

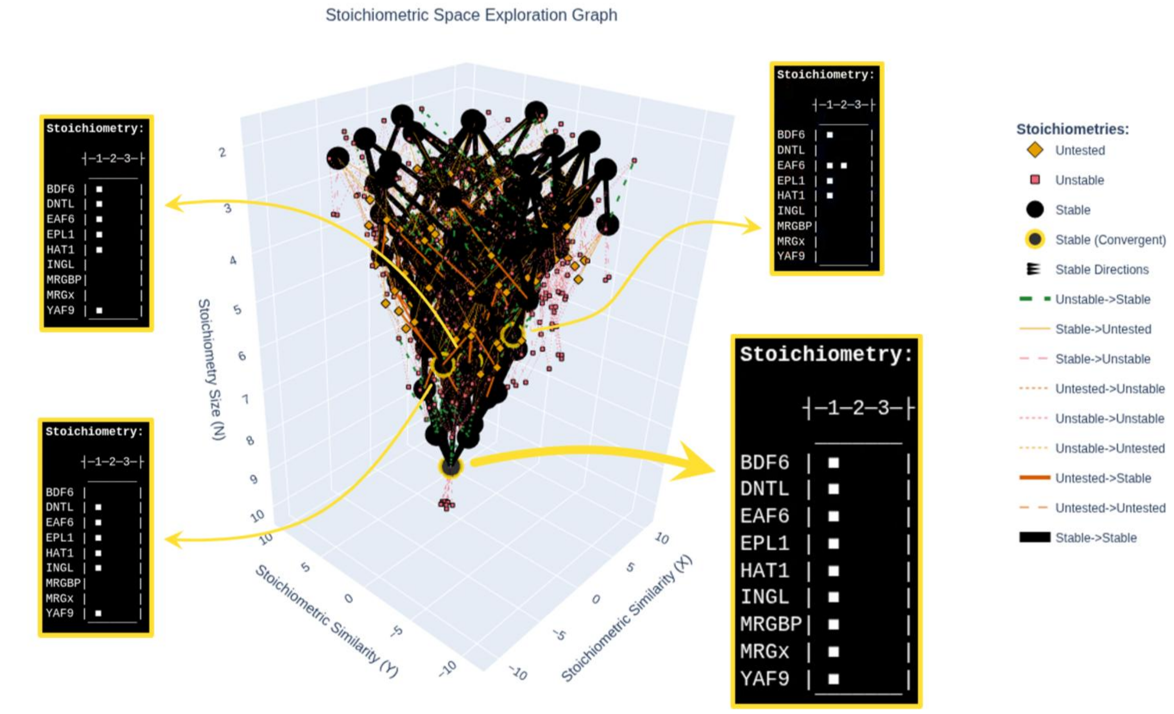


**Supplementary Fig. 13: Stoichiometric space exploration graph**. Output graph from MultimerMapper's stoichiometric space exploration algorithm for NuA4. Each marker represents a protein combination (or stoichiometry) predicted by AlphaFold3. Lines represent parent-child relationships between stoichiometries from different sizes. The stoichiometries of the detected convergent protein combinations are shown in the insets. The proposed stoichiometry of the complex is highlighted with a bigger inset.

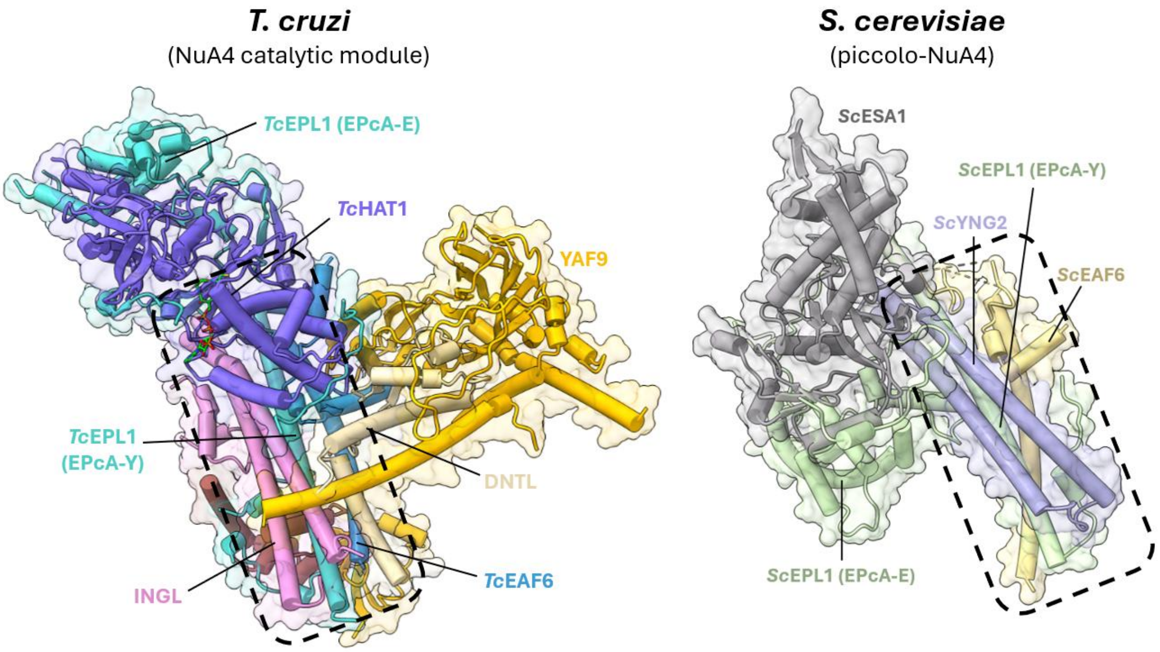


**Supplementary Fig. 14: Yeast piccolo-NuA4 module compared with the catalytic module of NuA4 from *T. cruzi*.** X-ray structure of piccolo-NuA4 from S. cerevisiae (PDB 5J9T) on the **right**, compared with the AF3 predicted structure of the catalytic module of NuA4 on the **left**. The boxes with dotted borders indicate the four alpha helices mentioned in the text.

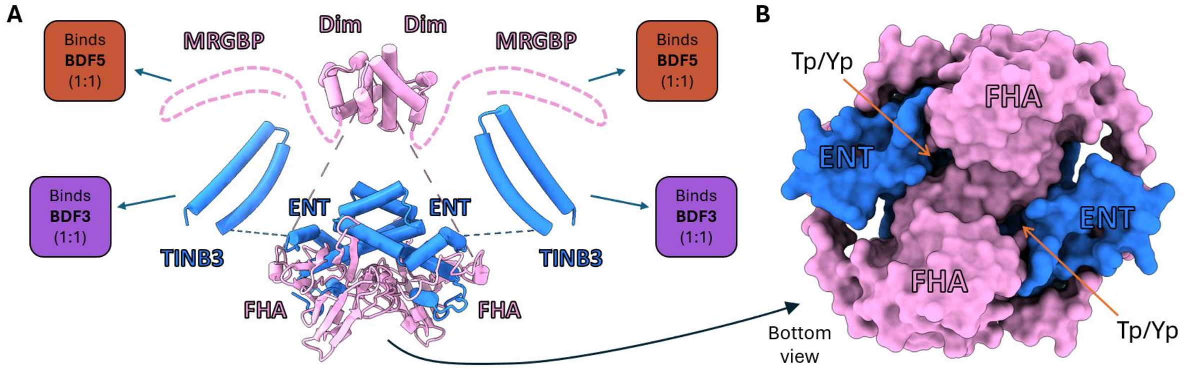


**Supplementary Fig. 15: Structure of the $2_{ENT}$:$2_{BDF5BP}$ heterotetramer predicted with AF3 and MultimerMapper**. (**A**) The heterotetramer exposes two MRG-binding domains (MRGBP, represented as dotted lines) and two BDF3 N-terminal α-helix-binding domains (TINB3), each with the potential to bind their partners with a 1:1 stoichiometric ratio. Segments with pLDDT less than 40 were removed for ease of visualization. (**B**) The structure formed by the FHA and ENT domains exposes phosphorylated threonine/tyrosine recognition pockets to the solvent. **Dim**: BDF5BP Dimerization domain. **TINB**: Trypanosomatid Interactor of N-terminal BET. **Tp**: Phosphorylated threonine. **Yp**: Phosphorylated tyrosine.

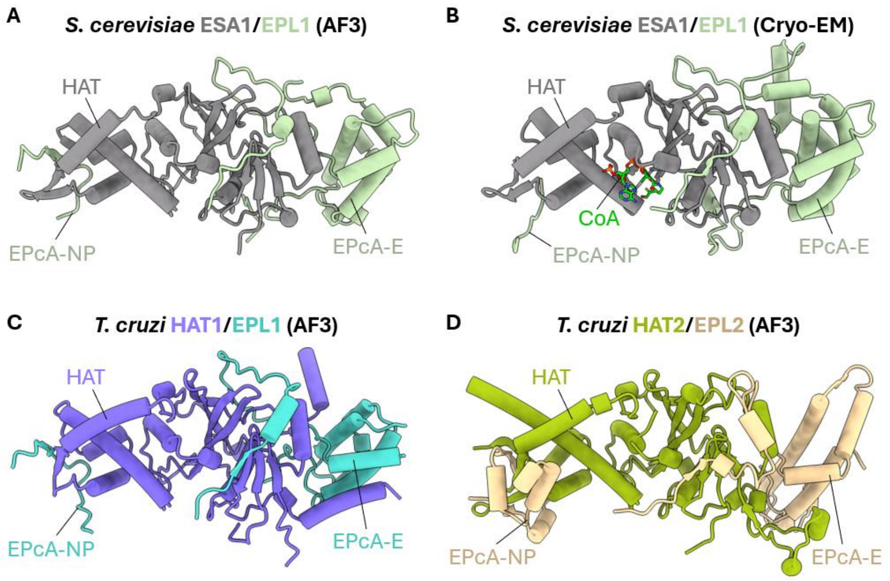


**Supplementary Fig. 16: Comparison between *S. cerevisiae* ESA1-ELP1, *T. cruzi* HAT1-EPL1 and HAT2-EPL2 heterodimers**. Predicted structure using AF3 server (**A**) and cryo-EM (PDB 7VVU) structure (**B**) of *S. cerevisiae* ESA1-EPL1 complex. Predicted structure using AF3 server of *T. cruzi* HAT1-EPL1 (**C**) and HAT2-EPL2 (**D**). Homologous domains are indicated. Disordered segments and domains not involved in the corresponding HAT-EPL interactions were removed to enhance visualization.

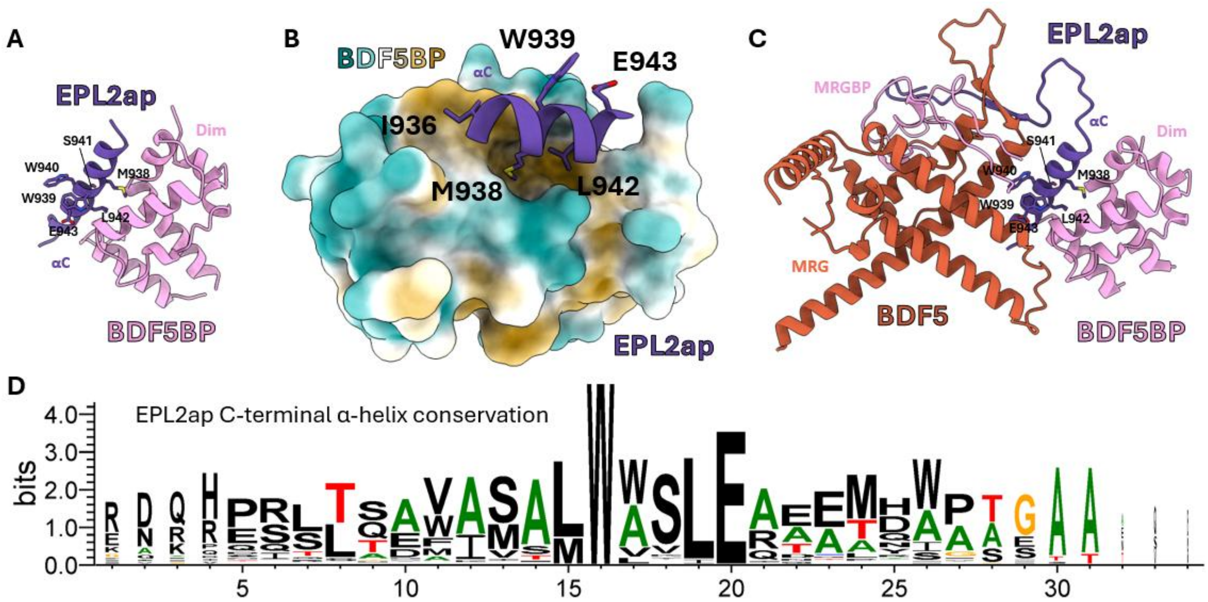


**Supplementary Fig. 17: Predicted interactions engaged by the C-terminal α-helix of EPL2ap**. (**A**) Predicted interaction between the C-terminal α-helix of EPL2ap and the Dim domain of BDF5BP from the AF3 heterodimeric prediction. (**B**) Close up of the binding region from panel A, showing solvent exposed surface of Dim, colored by hydrophobicity. (**C**) Predicted interaction between the C-terminal α-helix of EPL2ap, the Dim domain of BDF5BP and the MRG domain of BDF5 from the AF3 heterotrimer. (**D**) Logo representing the sequence conservation in bits of the C-terminal α-helix of EPL2ap across trypanosomatid species.

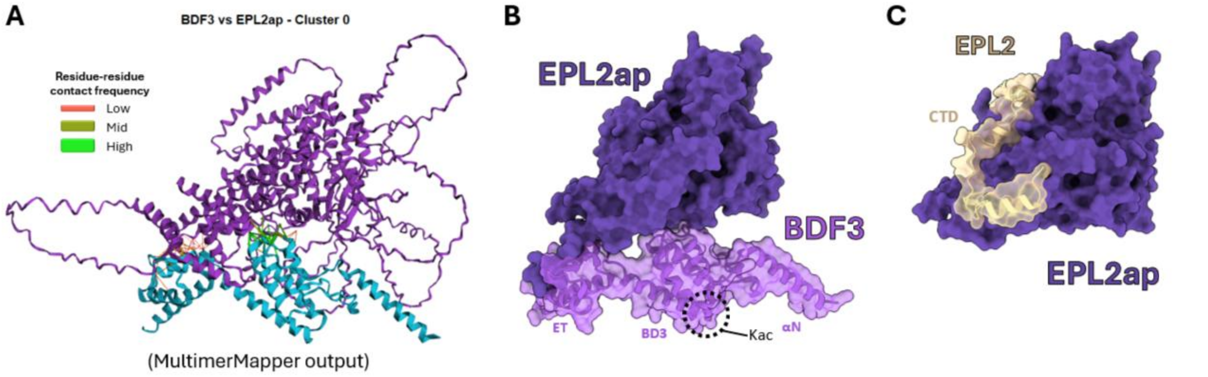


**Supplementary Fig. 18: Predicted interaction mode between EPL2ap and BDF3**. (**A**) Interactive output from MultimerMapper for the binding mode between EPL2ap and BDF3. Predicted residue-residue contacts are shown with lines, where red colors indicate lower frequency and green higher. (**B**) Representative structure of the binding mode from panel A explored with ChimeraX. Disordered loops were removed. (**C**) Representative structure of the binding mode between the CTD domain of EPL2 with EPL2ap explored with ChimeraX and aligned to the model from panel B. Notice that the surface involved are different.

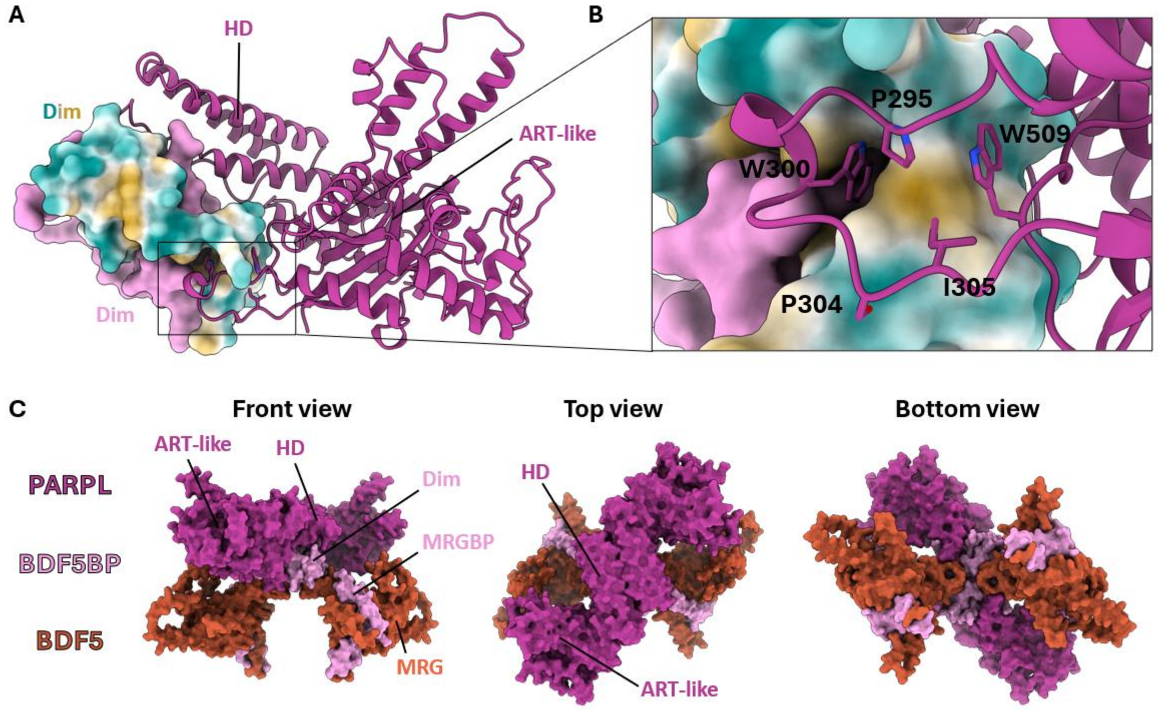


**Supplementary Fig. 19: Interaction between BDF5BP and PARPL**. (**A**) Interaction mode between the Dim domain of BDF5BP (two subunits) and PARPL (one subunit). (**B**) Close up of the contact region between the ART-like domain and Dim. (**C**) AF3 prediction using two subunits of each BDF5, BDF5BP and PARPL. Bromodomains, the FHA domain and disordered loops were removed to enhance visualization.

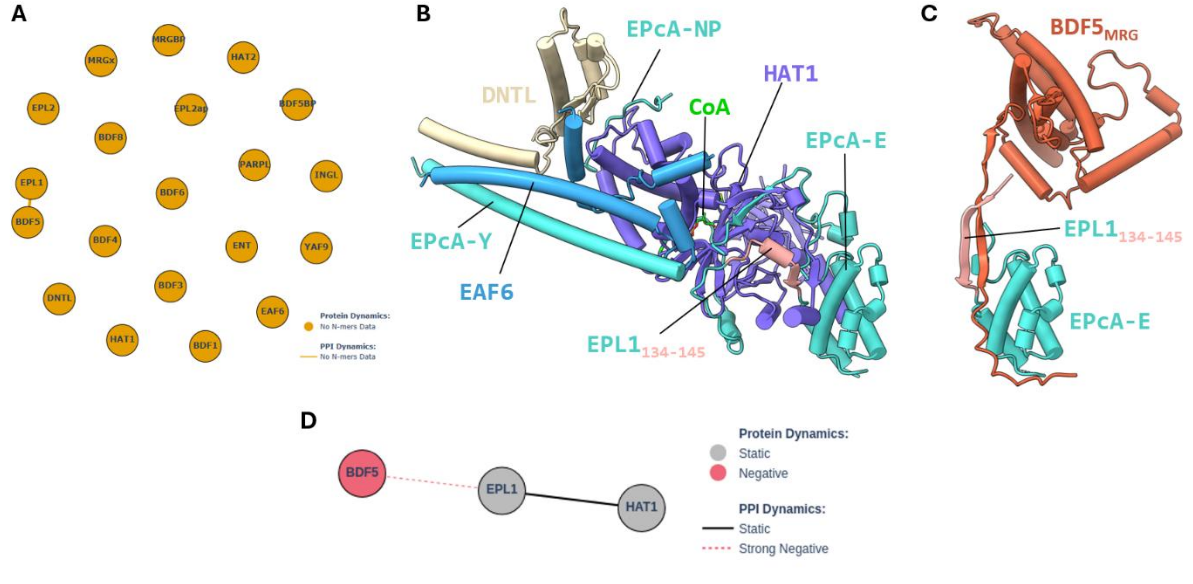


**Supplementary Fig. 20: Predicted crossed PPIs between components of NuA4 and CRKT**. (**A**) Predicted PPIs using MultimerMapper of crossed pairwise interactions between all the sequences from CRKT and NuA4. Only BDF5 and EPL1 were predicted to engage in a PPI. (**B**) Posterior view of NuA4 complex from **Error! Reference source not found.A**. The region of EPL1 involved in interactions with BDF5 is highlighted in pink (residues 134 to 145). (**C**) Predicted interaction mode between EPL1 and the MRG domain of BDF5. The structure is aligned with the model from panel B though the EPcA-E domain of EPL1. (**D**) PPI dynamics analysis of the BDF5-EPL1-HAT1 system using MultimerMapper.

### Supplementary Tables

**Supplementary Table 1: Fusion proteins used in each experimental round.**

| 3xHA-TurboID fusion | Abbreviature | Experimental round 1 | Experimental round 2 |
|---|---|---|---|
| Control without TurboID | NoTurbo | ✓ | ✗ |
| GFP | GFP | ✓ | ✓ |
| GFP-3xNLS | NLS | ✓ | ✓ |
| HA-BDF2 | BDF2 | ✓ | ✗ |
| HA-BDF3 | BDF3 | ✗ | ✓ |
| HA-BDF4 | BDF4 | ✗ | ✓ |
| HA-BDF5 | BDF5 | ✓ | ✗ |
| HA-BDF6 | BDF6 | ✓ | ✗ |
| BDF8 | BDF8 | ✓ | ✗ |

**Supplementary Table 2: Interacting proteins of each intersection in the Venn diagram.**

| Intersection | Total | Shared proteins |
|---|---|---|
| **BDF3 BDF4 BDF5 BDF8** | 3 | C4B63_2g92 C4B63_7g80 C4B63_10g308 |
| **BDF3 BDF4 BDF5** | 4 | C4B63_6g222 C4B63_89g65 C4B63_242g4 C4B63_28g81 |
| **BDF4 BDF5 BDF8** | 2 | C4B63_76g133 C4B63_1g51 |
| **BDF5 BDF6 BDF8** | 1 | C4B63_20g159 |
| **BDF3 BDF4** | 2 | C4B63_193g10 C4B63_12g154 |
| **BDF5 BDF6** | 4 | C4B63_4g143 C4B63_41g80 C4B63_8g485 C4B63_46g45 |
| **BDF6 BDF8** | 2 | C4B63_95g27 C4B63_18g1091c |
| **BDF3** | 7 | C4B63_20g132 C4B63_14g84 C4B63_40g26 C4B63_41g197 C4B63_70g92 C4B63_87g21 C4B63_2g321 |
| **BDF4** | 8 | C4B63_118g20 C4B63_63g19 C4B63_84g52 C4B63_25g318 C4B63_39g335 C4B63_7g431 C4B63_23g256 C4B63_42g92 |
| **BDF5** | 3 | C4B63_7g224 C4B63_58g53 C4B63_25g212 |
| **BDF6** | 9 | C4B63_10g400 C4B63_43g129 C4B63_30g209 C4B63_25g289 C4B63_89g383c C4B63_24g171 C4B63_16g124 C4B63_22g316 C4B63_425g10c |
| **BDF8** | 20 | C4B63_10g431 C4B63_275g16 C4B63_43g152 C4B63_51g192 C4B63_235g11 C4B63_92g53 C4B63_10g310 C4B63_7g208 C4B63_2g809 C4B63_137g1 C4B63_23g134 C4B63_9g493 C4B63_24g264 C4B63_83g31 C4B63_45g131 C4B63_75g68 C4B63_10g140 C4B63_2g7 C4B63_12g238 C4B63_7g270 |

**Supplementary Table 3: Oligonucleotides used for cloning.**

| Name | Sequence | Enzyme | Comment |
|---|---|---|---|
| **3xHA-TurboID-Fw** | AGTGATCGATTGGCCGCCACCATGTACCCG | ClaI | pTcTurboID cloning |
| **3xHA-TurboID-Rv** | AGTGATCGATTCGTCGGCCCTGCTGAATTCC | ClaI | pTcTurboID cloning |
| **GFP-Fw** | AAGGATCCATGAGTAAAGGAGAAGAACTTTTCACTGG | BamHI | pENTR3c cloning (GFP) |
| **GFP-Rv** | AACTCGAGTTTGTATAGTTCATCCATG | XhoI | pENTR3c cloning (GFP) |
| **BDF2-Fw** | AAAGGATCCATGTATCCGTATGATGTGCCGGATTATGCTGG GAAGCGTGGGCGT | BamHI | pENTR3c cloning (BDF2) HA tag (N-terminal) |
| **BDF2-Rv** | AAAGATATCTTATTCATCATCACTCTCATCATCATAATAAAAC TCTTC | EcoRV | pENTR3c cloning (BDF2) |
| **BDF3-Fw** | AAGGATCCATGGGCTCTACGGGTCGG | BamHI | pENTR3c cloning (BDF3) |
| **BDF3-Rv** | AACTCGAGCCTCGTCCTCCACCGCC | XhoI | pENTR3c cloning (BDF3) |
| **BDF4-Fw** | CGGTACCATGTATCCGTATGATGTGCCGGATTATGCTGCTG GTGGCTCTGTTCAA | KpnI | pENTR3c cloning (BDF4) HA tag (N-terminal) |
| **BDF4-Rv** | AAACTCGAGTCATTGCAAAAATTTCTCTATGCTCC | XhoI | pENTR3c cloning (BDF4) |
| **BDF5-Fw** | AAAGGATCCATGTATCCGTATGATGTGCCGGATTATGCTGA GGAGAGGCGGCAAAAG | BamHI | pENTR3c cloning (BDF5) HA tag (N-terminal) |
| **BDF5-Rv** | AAAGATATCCTACGATATTTCTTTTCCAATTTCCG | EcoRV | pENTR3c cloning (BDF5) |
| **BDF6-Fw** | AAAGGATCCATGTATCCGTATGATGTGCCGGATTATGCTCG GCGGGAAGATTTACTGC | BamHI | pENTR3c cloning (BDF6) |
| **BDF6-Rv** | AAAGATATCTCATGCACCACGCAAATGC | EcoRV | pENTR3c cloning (BDF6) |
| **BDF8-Fw** | AAAGTCGACAATGGATTTGATTTTTGCTGAGGGAG | SalI | pENTR3c cloning (BDF8) |
| **BDF8-Rv** | AAATCTAGATTATCACAACAGGCCCACCTCG | XbaI | pENTR3c cloning (BDF8) |
